# A structured RNA balances DEAD-box RNA helicase function in plant alternative splicing control

**DOI:** 10.64898/2026.01.23.701338

**Authors:** Rica Burgardt, Julia Bauer, Maren Reinhardt, Natalie Rupp, Christoph Engel, Sören Lukas Hellmann, Maximilian Sack, Zasha Weinberg, Andreas Wachter

## Abstract

Eukaryotic gene expression is a multi-layered process influenced by multiple factors. One of them is the secondary structure of precursor mRNAs that can impact various aspects of their processing including alternative splicing (AS). Here, we report the functional characterization of the conserved RNA structural element *DEAD* that is located in DEAD-box RNA helicase (DRH) genes from land plants and serves as a sensor for RNA helicase activity by controlling AS. In *Arabidopsis thaliana*, it is found in *DRH1* and its closest paralog, regulating usage of an alternative splice site as part of a negative feedback loop. Accordingly, opening of the structure shifts splicing towards non-coding variants, thereby balancing transcript and protein levels. Interestingly, the system is specific to DRH1 and its paralog and does not react to related helicases, which is at least partially conferred by the disordered and RGG/RG motif-containing C-terminus of DRH1. The importance of *DEAD* is underlined by the observation that releasing this attenuation mechanism causes massive changes in AS – mainly intron retention and exon skipping – and gene expression and results in a severe stress phenotype. Thus, *DEAD* provides a critical buffering mechanism to fine-tune helicase levels and their global impact on RNA structure-responsive gene expression.

## Introduction

The ability of RNA to form intramolecular base-pairs allows it to fold into a diverse spectrum of secondary structures with varying degrees of complexity. These structures are dynamic and react to various conditions, such as temperature, transcriptional speed, subcellular localization or interaction with RNA-binding proteins (RBPs) ^1–5^. In the past two decades, RNA structures have increasingly been recognized as important elements of eukaryotic gene regulation ^6,7^. Their modes of action are diverse and include amongst others control of transcription, alternative splicing (AS), translation, and RNA localization as well as stability ^8^. Transcriptome-wide structural profiling studies in both animals and plants revealed global patterns associated with efficient splicing, such as low structuredness of nucleotides directly upstream of splice sites ^2,9,10^. However, local structures are also frequently used to control individual AS events, e.g. by obstructing splice sites or other *cis*-regulatory elements or by bringing a particular set of splice sites into close proximity ^7^. Interestingly, up to 4% of AS events conserved among vertebrates were shown to contain RNA structures overlapping with at least one of the splice sites ^11^. The *Dscam1* gene from fruit fly, which encodes the impressive number of 38,016 possible AS isoforms, utilizes long-range base-pairing between a set of complementary sequences to select from a cluster of mutually exclusive exons ^12^. Specific interactions between structured RNAs and other molecules further expand the regulatory potential. Eukaryotic riboswitches from the fungal and plant kingdoms possess a structural element that recognizes the small metabolite thiamin pyrophosphate and thereby adopts a fold that exposes an alternative splice site and shifts the splicing pattern of the riboswitch-harboring gene towards unproductive transcripts ^13–15^. AS of human *MALT1* is regulated by RNA motifs that obstruct the splice sites of an alternative exon and are antagonistically modulated by two RBPs, thereby altering splice site accessibility ^16^.

RNA structures can be actively adjusted by RNA helicases, a large group of proteins with diverse functions. The majority of them can be grouped into two superfamilies; members of both share two highly conserved helicase domains that catalyze the separation of RNA strands. The largest subfamily is represented by DEAD-box RNA helicases which carry a conserved amino acid motif (DEAD, Asp-Glu-Ala-Asp) within their N-terminal helicase domain ^17^. Studies in yeast showed that DEAD-box helicases do not unwind long regions of double-stranded RNA, but instead act as local remodelers of secondary structure or RNA-protein complexes ^18^. They are involved in virtually every RNA-related process in the cell, ranging from transcription via splicing and RNA export to translation and RNA decay ^19^. In several instances, they were shown to affect AS via modulation of local stem-loop structures ^20–22^ and/or RNA-protein complexes ^23,24^. Plants evolved an expanded set of DEAD-box helicases: The model species *Arabidopsis thaliana* has 56 protein-coding genes of this family, compared to 37 in human ^17,25^.

In this study, we describe a novel plant-specific structured RNA element that employs AS-mediated feedback regulation to modulate RNA helicase activity. It was identified in a previous screen for conserved structured RNAs in the non-coding regions of plant genomes ^26^. The motif consists of a single stem-loop that occurs in DEAD-box RNA helicase genes of all land plant clades and was accordingly named *DEAD*. We could show that it acts as a sensor for the activity of *A. thaliana* DEAD-BOX RNA HELICASE 1 (DRH1) by obstructing an alternative splice site that is used for negative feedback control. Specific recognition of this particular helicase is at least in part conferred by its C-terminal RGG motif. DRH1 was identified as a major regulator of AS and gene expression with multiple stress-related target genes. Consequently, its overexpression resulted in a strong bleaching phenotype, illustrating the importance of this regulatory circuit for the plant. Our results highlight how a single *cis*-regulatory element can have a major impact on plant gene expression by controlling levels of a *trans*-acting factor.

## Results

In a previous study, several candidate RNA elements were identified in the non-coding regions of plant genomes that have a potentially conserved structure and overlap with alternative splice sites ^26^. One of the top hits from this search had already been described as a conserved intronic element in angiosperms ^27^, but no function had been assigned to it. Since it appears almost exclusively within genes encoding DEAD-box RNA helicases (Supplemental Data S1), we called it the *DEAD* motif.

Our previous search was limited to RefSeq 204 ^28^, which is heavily biased towards angiosperms. We therefore searched further genomes from the basal plant clades available in NCBI Datasets (www.ncbi.nlm.nih.gov/datasets) for similar sequences using nucleotide BLAST. We found 14 additional sequences in 9 species that fit the motif very well in both sequence and structure (Supplemental Data S1, Supplemental File 1), indicating that it was kept over long evolutionary distances. *DEAD* is not found in algae but is highly conserved among land plants, with hits in bryophytes, lycophytes, ferns, gymnosperms, and angiosperms (Fig. 1a). In total, we identified 353 high-confidence instances in 123 species ^26^ (Supplemental Data S1). The motif is predicted to form a single hairpin with a long stem of 19-20 nt and a short terminal loop encompassing 3-5 nt (Fig. 1b). Sack et al. also identified a conserved alternative 3’ splice site within the motif and confirmed its usage in several species ^26^. We therefore wanted to investigate if the structure is in fact involved in AS control. As our model organism, we chose *A. thaliana*, where the motif appears in two genes: *DEAD-BOX RNA HELICASE 1* (*DRH1*, also called *RH14*) and its paralog, *RNA HELICASE 46* (*RH46*).

**Figure 1:**
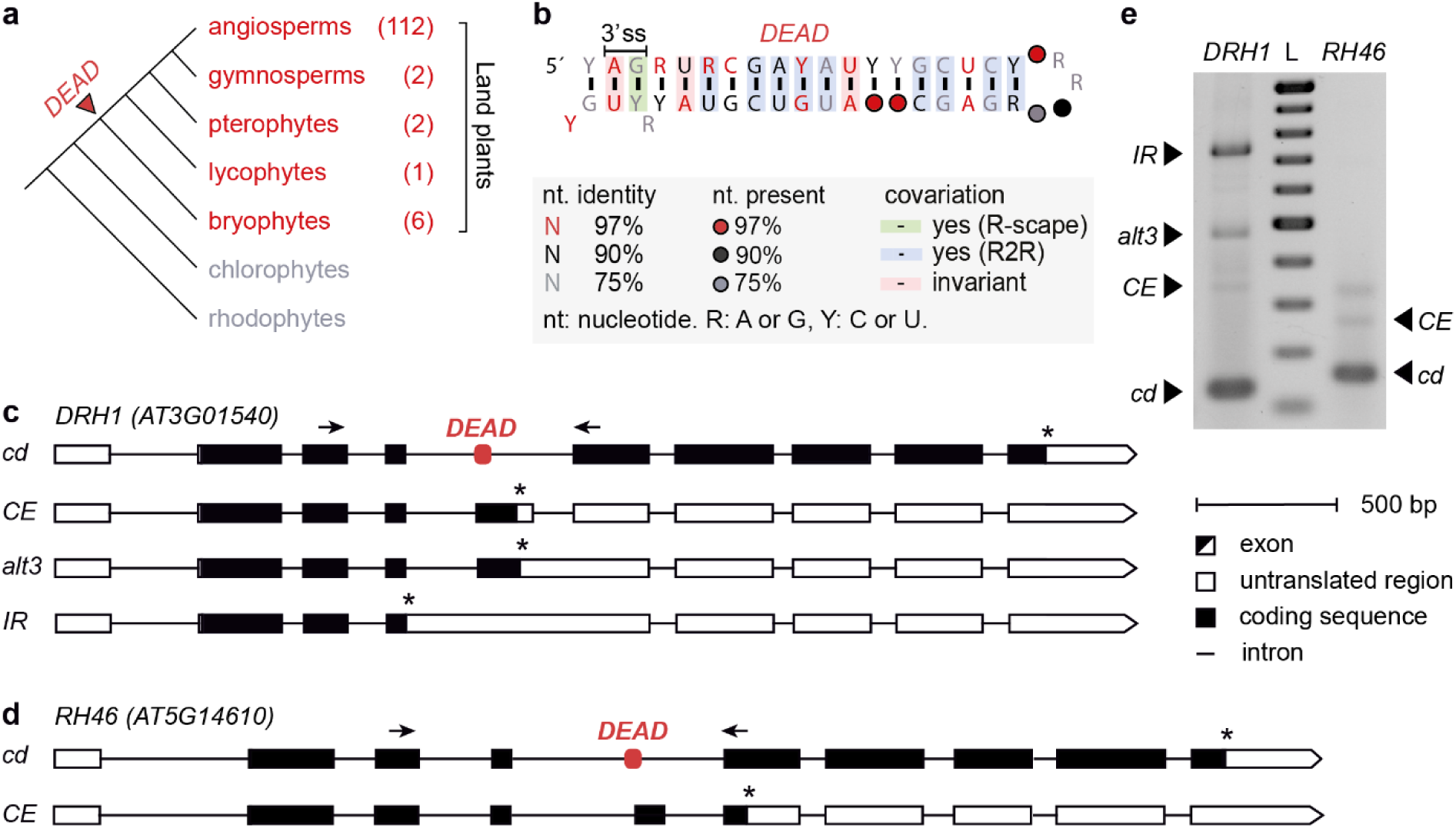
The *DEAD* motif is conserved among land plants. **a:** Schematic representation of plant evolution; clades carrying the *DEAD* motif are marked in red. Numbers in parentheses indicate the number of species per clade that contain at least one instance of the motif. The red arrowhead marks the point of *DEAD* emergence. **b:** Consensus sequence and predicted structure of the *DEAD* motif across all species it was found in. Shading indicates covariation (green: significant covariation as determined by R-scape, blue: covariation determined by R2R without statistical analysis, red: invariant). 3’ ss: alternative 3’ splice site. Figure is based on the *DEAD* motif from Sack et al. (2025)^26^ that we updated with additional species from the bryophyte, pterophyte, and gymnosperm clades. **c-d:** Gene models of *DRH1* (**c**) and *RH46* (**d**) showing AS variants involving the intron that *DEAD* is located in. Exons, introns, coding sequence and untranslated regions are represented by boxes, lines, black and white shading, respectively. Arrows represent primers used for co-amplification RT-PCR in **e**; asterisks indicate stop codons. **e:** AS of *DRH1* and *RH46* in *A. thaliana* Col-0 seedlings, RT-PCR bands representing the transcripts shown in **c** and **d** are marked with black arrowheads. The band above *RH46 CE* probably is a running artefact, as it does not appear in Bioanalyzer runs. L: Ladder, from 100 bp upwards in 100 bp increments.

### *DRH1* is autoregulated via alternative splicing

We first set out to characterize the AS landscape derived from the two helicase genes. In both cases, the motif is located within a long intron and overlaps with an alternative 3’ splice site as mentioned above (Fig. 1c-d). For *DRH1*, there are four transcript isoforms involving the intron that harbors *DEAD* (Fig. 1c). Complete removal of the intron results in a protein-coding transcript (*cd*); use of the alternative 3’ splice site within *DEAD* (Fig. 1b) can lead to either inclusion of a cassette exon (*CE*) or a variant retaining the 3’ end of the intron (*alt3*); and lastly, the intron can be completely retained (*IR*). In case of *RH46*, corresponding coding (*cd*) and cassette exon (*CE*) variants could be identified (Fig. 1d). For both genes, the main splicing product detected by RT-PCR is *cd* (Fig. 1e). Apart from *cd*, all transcripts contain premature termination codons (PTCs) and are therefore expected to be non-coding (*nc*) and probably targeted by RNA surveillance systems. Accordingly, the *alt3* variant was shown to be degraded via nonsense-mediated decay ^26^.

Since autoregulation via splicing to non-coding transcripts is a common mechanism for RBPs ^29,30^, we tested if DRH1 can influence AS of its own pre-mRNA. We constructed a splicing reporter for *DRH1*, consisting of the 5’ sequence of the gene until shortly after the alternatively spliced intron with an attached *GFP* (Fig. 2a). This reporter was transformed transiently into *Nicotiana benthamiana* together with a construct for expression of the *DRH1* coding sequence (CDS) or a control protein (Luciferase, LUC). Notably, there are two protein isoforms of DRH1 that differ only in the presence (DRH1-I) or absence (DRH1-II) of a serine close to the C-terminus, caused by a NAGNAG site at the 3’ end of the last intron. Both isoforms are capable of autoregulation via AS: When LUC was co-transformed, only the coding variant was detectable, whereas co-transformation of DRH1 led to increased splicing towards the *alt3* and *IR* variants (Fig. 2b). We also measured this effect on protein level: As all non-coding variants contain a PTC, GFP fluorescence directly correlates with the amount of *cd* transcript. Reporter fluorescence was reduced to ∼30% in presence of DRH1 when compared to LUC samples (Fig. 2c), confirming that DRH1 negatively auto-regulates its level. We could not find a difference between the two protein isoforms regarding their regulatory function. In the following and if not specified otherwise, figures show the data for the longer isoform DRH1-I; the data for DRH1-II can be found in the supplement.

**Figure 2:**
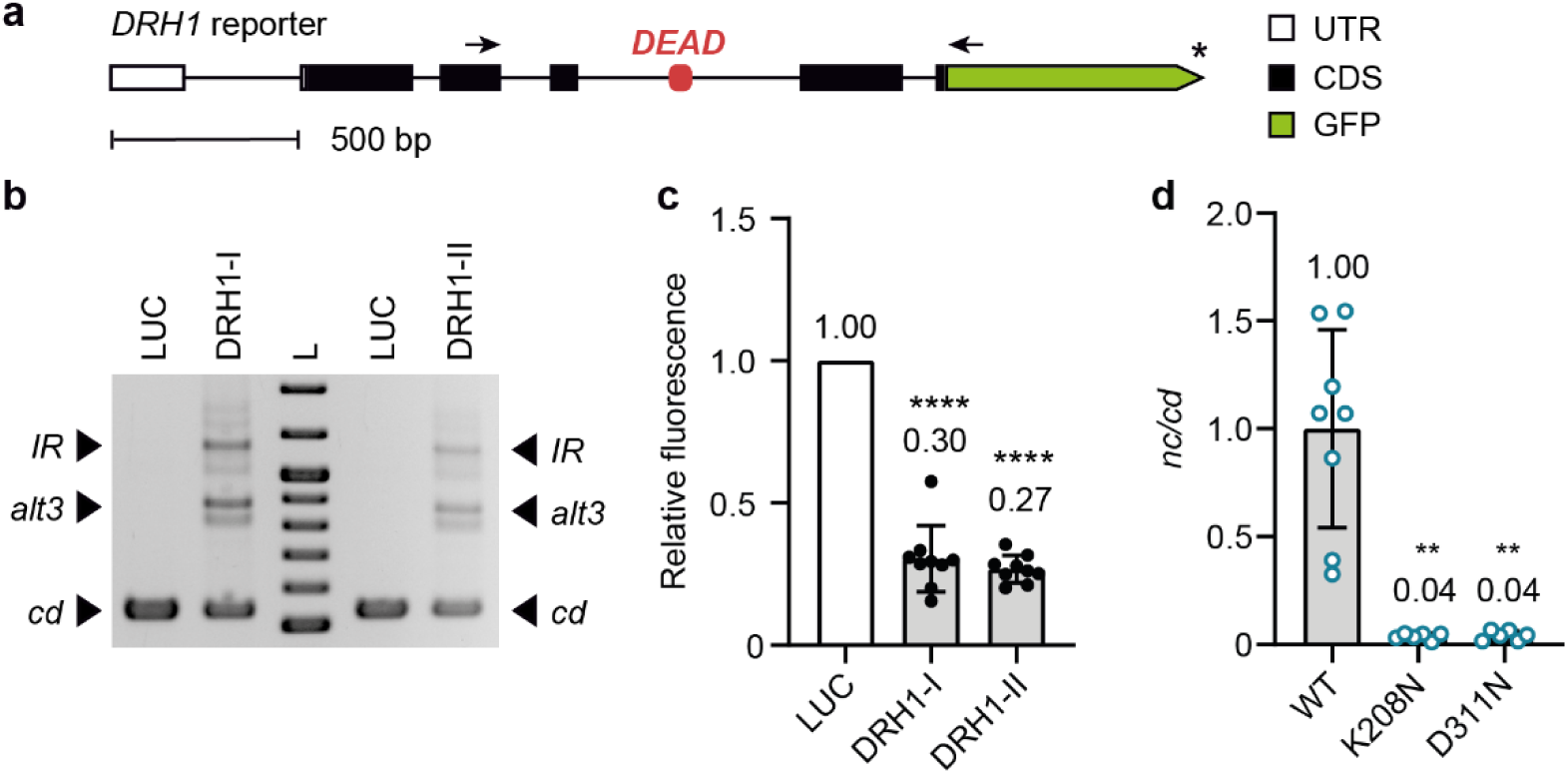
*DRH1* autoregulation via AS requires helicase activity in *N. benthamiana*. **a:** Gene model of the *DRH1* splicing reporter. Exons, introns, CDS and UTRs are represented by boxes, lines, black and white shading, respectively; *GFP* tag is depicted in green. Arrowheads represent primers used for co-amplification RT-PCR in **b** and **d**; asterisk indicates stop codon. **b:** Reporter splicing in *N. benthamiana* when co-transformed with constructs for the two DRH1 protein isoforms or a control protein (Luciferase, LUC). Transcript variants are marked with black arrowheads, the double bands for *alt3* and *IR* correspond to a single peak in Bioanalyzer runs and therefore probably represent running artefacts. L: ladder, from bottom to top: 0.5-1 kb in 0.1 kb increments, 1.2 kb, 1.5 kb. **c**: DRH1-mediated decrease of reporter fluorescence upon transient expression in *N. benthamiana*. Fluorescence in presence of LUC was set to 1. *n* = 9. **d:** Bioanalyzer quantification of reporter AS in *N. benthamiana* upon co-transformation of WT DRH1-I or two DRH1-I mutants impaired in helicase activity. AS ratio in presence of WT DRH1-I was set to 1. *n* = 8 for WT, *n* = 6 for K208N and D311N. Mean values, standard deviations; individual data points represent biological replicates and are displayed as black or blue dots. Asterisks indicate significant change compared to co-transformation of LUC (**c**, one-sample t-test) or WT DRH1 (**d**, Kruskal-Wallis test followed by Dunn’s multiple comparisons test). **p < 0.01, ****p < 0.0001.

### AS-mediated autoregulation requires DRH1 helicase activity

Given that DRH1 is an RNA helicase, we hypothesized that it might autoregulate its own mRNA levels by unwinding the *DEAD* motif, which would expose the alternative 3’ splice site and therefore cause increased splicing towards the non-coding variants. Consequently, we tested if a DRH1 mutant lacking helicase activity also loses the ability to control AS. For the closest human homolog of DRH1, DDX5, several amino acids are known to be crucial for helicase activity ^31^. We chose to test two established mutants: *Hs*-DDX5-K144N, which is reported to have no helicase activity and shows reduced RNA binding, and *Hs*-DDX5-D248N, which retains partial ability to hydrolyze ATP and normal affinity to RNA ^32,33^. In both cases, the mutations are located in highly conserved motifs within the helicase domain and thus likely to have similar effects when applied to orthologs from other organisms. Mutation of the corresponding positions in DRH1 resulted in *At*-DRH1-K208N and *At*-DRH1-D311N which were tested in *N. benthamiana* using our *DRH1* splicing reporter. Both mutants induced a much weaker AS shift than the WT DRH1 CDS, with the *nc/cd* ratio dropping to ∼5% (Fig. 2d, SFig. 1a-e). This suggests that helicase activity of DRH1 is indeed required to control AS of its own mRNA. However, both mutants can still inhibit fluorescence of the *DRH1* reporter (SFig. 1f); so either the remaining AS shift is sufficient to repress protein expression, or there is a splicing-independent mechanism of autoregulation that does not require helicase activity. It has to be considered that impairment of helicase activity can also affect RNA binding/dissociation ^32,33^, which might at least partially explain the splice shift. It is therefore likely that DRH1 helicase activity is necessary for efficient usage of the alternative 3’ splice site, but the role of the structure required a direct assessment.

### The secondary structure of *DEAD* is involved in AS regulation

We next wanted to know if the predicted stem-loop structure of *DEAD* is required for autoregulation of *DRH1*. To this end, we created mutations of the motif both in the reporter context and in the full-length gene (Fig. 3a). Specifically, we introduced disruptive mutations (*DM1-3*) of varying length to impair base-pairing within the stem and thereby expose the alternative splice site. Note that only nucleotides within the bottom 10 bp of the stem were mutated, the top half was left intact. To make sure that any changes observed for these mutations are due to the change in structure, we also created compensatory mutations (*CM1-3*) designed to restore the hairpin, but with an altered sequence. If RNA folding is the only decisive factor for the motif’s function, the compensatory mutants should behave similar to the wild-type sequence whereas the disruptive mutations should result in a distinct AS pattern. Analysis of reporter splicing in stably transformed *A. thaliana* lines revealed that the WT version is spliced almost completely to *cd*, indicating that DRH1 activity is limited (Fig. 3b). All three disruptive mutations, however, caused a massive shift in the splicing pattern towards *nc* variants, particularly *alt3* (Fig. 3b-c, SFig. 2a-b). Interestingly, this shift was in a similar range for *DM2* and *DM3*, although the former opens a larger part of the stem (Fig. 3a). *DM3* is predicted to cause a loop within the stem which might lead to destabilization of the flanking sequences, thereby opening an even larger part of the structure. It is also notable that the *IR* variant increased as well, although it does not require the alternative splice site exposed by the mutation (SFig. 2b). This suggests that *DEAD* can also influence the use of distant splice sites, possibly by folding back over them or by recruitment of unproductive splicing complexes. Importantly, the observed effects could be compensated by restoring the predicted hairpin, supporting the conclusion that this structure is indeed formed *in vivo* and involved in AS regulation. All compensatory mutants shifted back to an AS pattern closely resembling that of the WT reporter (Fig. 3b-c, SFig. 2a-b). *CM1* and *CM2* exhibited slight overcompensation, as *alt3* and *IR* levels were reduced in comparison to WT. This could be explained by the missing bulge nucleotide opposite the alternative splice site which probably renders the WT conformation of the stem less stable (Fig. 3a). To see if DRH1 is required for usage of the alternative splice site, we also tested these reporters transiently in *N. benthamiana*, where DRH1 levels can be controlled (SFig. 2c-e). Here, disruption of the *DEAD* structure led to only minor changes in splicing when DRH1 was present and had no effect in its absence, indicating that opening of the stem alone is not sufficient to cause increased use of the alternative 3’ splice site (SFig. 2c). DRH1 might therefore not only be needed to unwind the motif, but also to perform other functions, possibly recruitment of additional splicing regulators. Taken together, the structure of *DEAD* plays an important role in controlling transcript levels of *DRH1* via AS, probably by obstructing the alternative 3’ splice site.

**Figure 3:**
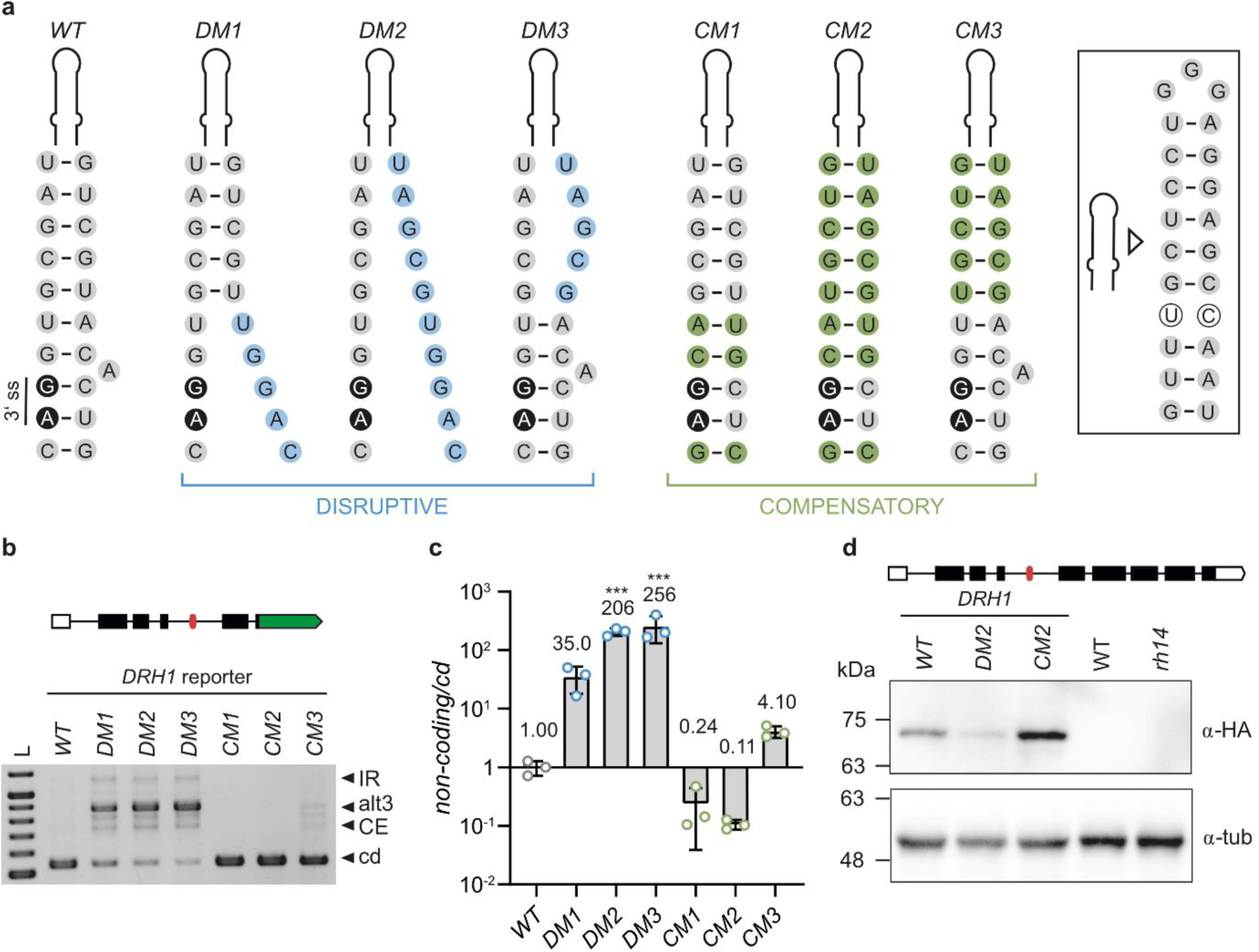
The *DEAD* motif is involved in *DRH1* AS regulation in a structure-dependent manner. **a:** Schematic representation of the base pairing potential of *DEAD* in its wild-type form as well as disruptive and compensatory mutations. Only the bottom 10 nt were mutated. Alternative 3’ splice site (3’ ss) is marked in black, mutated nucleotides are shaded in blue or green. Sequence and pairing potential of upper part of hairpin is shown in box, with mismatching nucleotides represented by open circles. **b:** Representative gel image of reporter AS in *A. thaliana*. AS variants are indicated by black arrowheads, the double bands observed for the *alt3* and *IR* transcripts correspond to a single peak in Bioanalyzer runs and therefore represent running artefacts. L: ladder, bottom to top: 0.5-1 kb in 0.1 kb increments, 1.2 kb. **c:** Bioanalyzer quantification of WT and *DEAD* mutant reporter AS in stably transformed *A. thaliana* lines, AS ratio of WT reporter was set to 1. *n* = 3. Mean values, standard deviations; individual data points represent independent lines and are depicted as colored dots. Asterisks indicate significant change compared to the WT reporter (one-way ANOVA followed by Dunnett’s multiple comparisons test, ***p < 0.001). **d:** Western Blot of *A. thaliana* lines carrying *pUBQ:DRH1* genomic constructs with WT or mutant *DEAD* motif in *rh14* background. Detection of HA-tagged DRH1 and tubulin (tub) as loading control are shown.

To see if the changes in *DRH1* AS also result in altered DRH1 protein levels, we introduced the *DRH1* full-length gene carrying either WT or mutated versions of *DEAD* into the previously described T-DNA insertion line *rh14* ^34,35^ under control of the *UBQ10* promoter. The AS patterns from these constructs mimicked that of the WT and mutant reporters, indicating that *DEAD* fulfils the same function also in its full-length genetic context (SFig. 2f). Western Blot analysis showed that there is indeed less protein detectable when the bottom half of *DEAD* is opened compared to the two closed-stem conformations (Fig. 3d, SFig. 2i-j). In addition, we looked at the splicing pattern of endogenous *DRH1* and observed a shift towards the non-coding variants for the WT and *CM2* conformations of *DEAD* (SFig. 2g), probably due to the elevated *DRH1* expression in these lines resulting from use of the *UBQ10* promoter (SFig. 2h). In the *DM2* mutants, this shift was less pronounced, suggesting a lower amount of functional DRH1 protein (SFig. 2g). These findings demonstrated that *DEAD* can control not only transcript, but also protein levels of DRH1.

### RGG motif confers functional specificity to DRH1 and RH46

The DEAD-box RNA helicase family in *A. thaliana* contains 56 protein-coding members ^25^, only two of which carry the *DEAD* motif. We wanted to see if there is any specificity in its recognition. Accordingly, we analyzed AS of the *DRH1* splicing reporter in *N. benthamiana* in presence of its closest homologs, namely RH46, RH30, and RH20. As *RH46* contains the same NAGNAG site as *DRH1*, it also features two isoforms distinguishable by the presence of a serine in the C-terminus. In addition, we intended to test RH40, another closely related helicase, but failed to obtain functional clones due to apparent cytotoxicity even of intronized CDS constructs in bacteria. Instead, we included UPF1, which is also an RNA helicase and was shown to interact with DRH1 ^34,36^. Of these proteins, only RH46 was able to impact AS of *DRH1* in a similar manner as DRH1 itself (Fig. 4a). In presence of RH20 and UPF1 only the coding variant was detectable, whereas co-expression of RH30 led to very small amounts of non-coding transcripts (Fig. 4a-b, SFig. 3a-b). On fluorescence level, we found that co-expression of all helicases affected the *DRH1* reporter output, although RH46 was again the only one with a similarly strong effect as DRH1 itself (Fig. 4c). Regarding AS control of *DRH1*, RH46 even seemed to exceed the impact of its paralog (Fig. 4b, SFig. 3a-b); however, it also gave a stronger signal in Western blot analysis which might account for the difference (Fig. 4d). Taken together, DRH1 levels might be regulated via splicing-independent mechanisms by all the tested helicases. However, the main response to opening of the *DEAD* stem, increased splicing towards *alt3* (Fig. 3b), was only observed upon co-expression of DRH1 or its paralog RH46. This suggests that *DEAD* is either not recognized as a substrate for unwinding by the other helicases, or they might unwind the hairpin, but lack another function that is required to induce an AS response.

**Figure 4:**
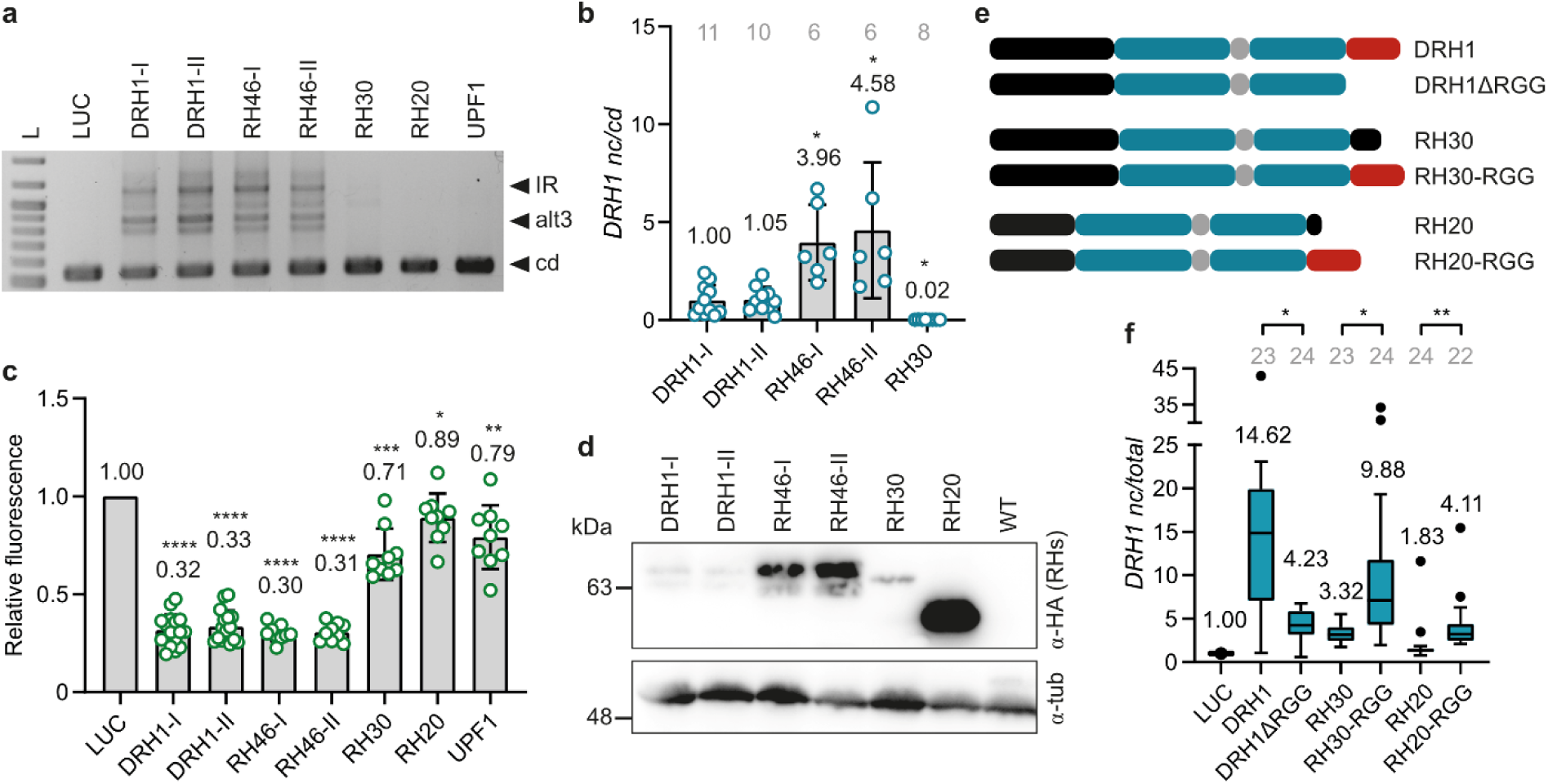
RH46 and RH30 can crossregulate *DRH1* via AS in *N. benthamiana*. **a:** AS response of *DRH1* reporter upon co-expression of different helicases in *N. benthamiana*. Different variants are indicated by arrowheads; the apparent double bands are running artefacts. L: Ladder, 0.5-1 kb in 0.1 kb increments, 1.2 kb, 1.5 kb. **b:** Bioanalyzer quantification of *DRH1* reporter AS on transcript level upon transient co-expression of DRH1, RH46, and RH30. AS ratio in presence of DRH1-I was set to 1, *n* is given in grey letters above each bar. **c:** Fluorescence inhibition of the *DRH1* reporter by different helicases upon transient expression in *N. benthamiana*. Fluorescence in presence of a control protein (Luciferase, LUC) was set to 1. *n* = 16 for DRH1-I and DRH1-II, *n* = 9 for all others. Mean values, standard deviations; individual data points represent biological replicates and are depicted as colored dots. Asterisks indicate significant change compared to DRH1 (**b**, Kruskal-Wallis test followed by Dunn’s multiple comparisons test) or LUC (**c**, one-sample t-test) co-transformation. **d:** Western Blot showing transient expression of different helicases in *N. benthamiana*. Each sample represents a pool from 5 transformed leaves. **e:** Model of WT and artificial helicase constructs. N- and C-termini, helicase domains, and linkers are shown as black, blue, and grey boxes, respectively; RGG motif is depicted in red. **f:** qPCR quantification of *DRH1* reporter AS upon transient co-expression of different helicases with and without RGG box. Transcript ratio in presence of LUC was set to 1, *n* is given in grey letters above each bar. Boxes, line, whiskers, and dots show 1^st^ to 3^rd^ quartile, median, data range up to 1.5 x interquartile range, and outliers, respectively. Numbers represent the mean. Significant changes between constructs with and without RGG box are depicted by asterisks above brackets (Kruskal-Wallis test followed by Dunn’s multiple comparisons test). *p < 0.05, **p < 0.01, ***p < 0.001, ****p < 0.0001.

Interestingly, DRH1 possesses a C-terminal RGG/RG motif ^37^ that is lacking in RH20 and is much shorter in RH30. We wondered if this disordered region might confer functional specificity among the helicases. To test this, we on one hand deleted the C-terminus from DRH1 and on the other hand fused it to RH20 and RH30 in place of their natural C-terminus (Fig. 4e). Splicing analyses in *N. benthamiana* revealed that the RGG/RG motif is indeed involved in *DRH1* AS control: Presence of the domain caused a ∼2- to 3-fold stronger shift towards the non-coding transcripts, regardless of the helicase it was attached to (Fig. 4f). As the C-terminus of RH46 looks similar to that of DRH1 (SFig. 4), it is likely that the RGG/RG motif at least contributes to their specific control of *DEAD*-mediated AS.

### DRH1 and RH46 are part of regulatory feedback loops

The specific role of DRH1 and RH46 suggested a negative feedback control of the two paralogs, prompting us to analyze knockout lines. As the T-DNA insertion in *rh14* is relatively close to the 3’ end of the gene and therefore might not have a functional impact, we identified the premature stop codon via 3’ RACE and cloned the resulting transcript (SFig. 5a, c). The truncated protein potentially translated from it has almost no regulatory activity on *DRH1* AS in *N. benthamiana* (Fig. 5a-c), so regarding autoregulation, *rh14* can probably be classified as a knockout. We also generated an *rh46* mutant via CRISPR-Cas9 mutagenesis (SFig. 5b) and crossed it with *rh14* to obtain an *rh14rh46* double knockout. Except for a mild delay in the onset of flowering for the double mutant (Fig. 5d), we observed no phenotypical changes, indicating that DRH1 and RH46 are dispensable under ideal growth conditions. To determine the impact of the two helicases on AS regulation of their respective pre-mRNAs, we quantified transcript levels and AS patterns of *DRH1* and *RH46* in the three mutants. As a consequence of the T-DNA insertion, total transcript levels of *DRH1* decrease to ∼10% in *rh14* and the double mutant compared to WT seedlings, and AS shifts strongly with a decrease of *nc*, confirming the AS-mediated autoregulation (Fig. 5e-f, SFig. 5d-f). In *rh46* mutants, there is no substantial change in either transcript levels or AS pattern of *DRH1*, indicating that RH46 has no great regulatory influence on its paralog. This is not true in the reverse: *RH46* total transcript slightly increases in the *rh14* line, indicating that DRH1 represses its paralog (Fig. 5g). Accordingly, the AS pattern of *RH46* shifts with a relative decrease of the non-coding *CE* transcript in *rh14* (Fig. 5h, SFig. 5g-i), confirming that AS takes part in this repression. In *rh46* plants, *RH46* shows a strong AS shift towards the *CE* variant (Fig. 5h) caused entirely by decrease of the *cd* transcript (SFig. 5g). *RH46 CE* levels remain unchanged in *rh46* plants (SFig. 5h), indicating that RH46 has no major autoregulatory role. Taken together, DRH1 represses itself and *RH46* via AS, but not vice versa. Considering that *RH46* is the more weakly expressed gene in virtually every tissue and developmental stage, based on publicly available expression maps ^38–40^ and our own RNAseq data (see later sections, SFig. 6), DRH1 might repress *RH46* globally, or low *RH46* expression caused by another factor results in its missing impact on *DRH1* AS.

**Figure 5:**
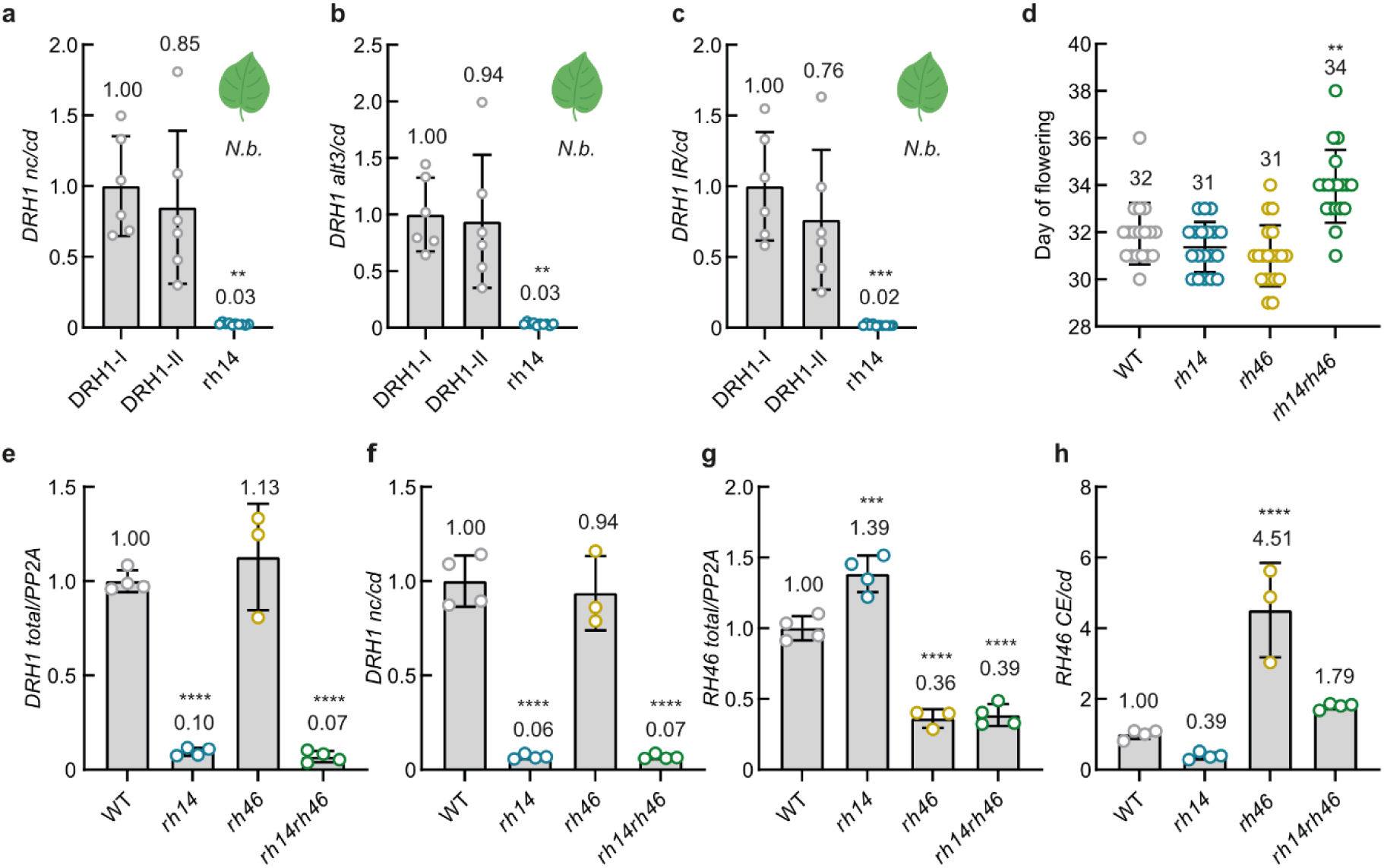
AS of *DEAD*-containing transcripts in *A. thaliana* is determined by DRH1. **a-c:** Bioanalyzer quantification of WT *DRH1* reporter AS in *N. benthamiana* (*N.b.*) upon co-transformation of WT DRH1 (both isoforms) or the protein variant potentially expressed in the *rh14* mutant. AS ratio of WT DRH1-I is set to 1. *n* = 6 for DRH1-I and DRH1-II, *n* = 10 for rh14. **d:** Flowering time (days after sowing) of *A. thaliana* WT plants or different mutants. Recorded was the day on which the inflorescence reached 1 cm in length. n = 18 for WT, 19 for *rh14* and *rh14rh46*, and 20 for *rh46*. **e-h:** Quantitative PCR of *DRH1* (**e-f**) and *RH46* (**g-h**) transcripts in *A. thaliana* seedlings of different backgrounds. Transcript ratio of WT seedlings is set to 1. *n* = 3 for *rh46*, *n* = 4 for all others. Mean values, standard deviations; individual data points represent biological replicates and are depicted as colored dots. Asterisks indicate significant change compared to DRH1-I (**a-b:** Brown-Forsythe and Welch ANOVA followed by Dunnett’s T3 multiple comparisons test, **c:** Kruskal-Wallis test followed by Dunn’s multiple comparisons test) or WT background (**d:** Kruskal-Wallis test followed by Dunn’s multiple comparisons test, **e-h:** one-way ANOVA followed by Dunnett’s multiple comparisons test). **p < 0.01, ***p < 0.001, ****p < 0.0001.

To analyze possible functions of DRH1 and RH46, we performed whole-transcriptome sequencing of all knockout mutants and two lines expressing the *DRH1* genomic construct in *rh14* background (*pUBQ:DRH1*). However, only a small number of genes was differentially expressed (DE) or showed differential alternative splicing (DAS) in the mutants compared to WT seedlings (SFig. 7, Supplemental Data S2). Loss of *DRH1* and *RH46* resulted in 9 and 2 DAS genes, respectively; knockout of both paralogs increased this number to 20 (SFig. 7b-c, e). This suggests that the two helicases have a limited impact under our growth conditions, possibly due to redundancy with other members of the large DEAD-box RNA helicase family that may compensate for the loss of DRH1 and/or RH46.

### Elevated DRH1 expression is detrimental

As *DEAD* is limiting DRH1 expression, we wondered what would happen if this control is released. We therefore wanted to analyze a mutant overexpressing DRH1. Constitutive overexpression of CDS constructs did not work, as we could only obtain lines with wildtype-like transcript levels of *DRH1* (SFig. 8a-b). Accordingly, we went for an estradiol-inducible expression system ^41^. Seedlings carrying a CDS construct of *DRH1-I* or *DRH1-II* under control of the estradiol-inducible promoter were treated with either β-estradiol or a mock solution and incubated for up to 4 days. Expression of the transgenic mRNA reached its peak between 3 to 6 h after estradiol treatment and was almost down to background levels after 2 days (Fig. 6a). Maximum *DRH1* transcript levels varied among different lines and ranged between 11- and 77-fold overexpression 6 h after induction in the estradiol-treated samples compared to the mock control. In WT seedlings, the treatment did not elicit a response. DRH1 protein expression was already detectable 3 h after treatment but reached its peak after 24 h and stayed high until the 2-day time point (Fig. 6b). The AS pattern of endogenous *DRH1* changed accordingly, with a 2- to 3-fold shift towards the non-coding variants 6 h after induction (Fig. 6c). This again confirms that DRH1 is autoregulating its own transcript via AS in *A. thaliana*. Total levels of endogenous *DRH1* mRNA showed a downward trend upon DRH1 induction, probably because of the AS shift and degradation of the *alt3* transcript via NMD. Interestingly, the estradiol-treated plants developed a severe stress phenotype starting ∼2-3 days after treatment and progressing: Leaves bleached out, and the seedlings stopped growing (Fig. 6d, SFig. 8c), being in line with our observation that we could not obtain constitutive overexpression lines. As an indicator for plant fitness, we measured the electron transport rate (ETR) 4 days after DRH1 induction. ETR has previously been shown to correlate with net photosynthetic rates and to decrease under stress ^42–44^. Whereas ETR reached ∼30-40 µmol e^-^ m^-2^ s^-1^ in the mock samples and also in WT seedlings treated with estradiol, DRH1 overexpression lines showed a strongly reduced ETR upon estradiol induction (Fig. 6e, SFig. 8d). Therefore, although low DRH1 levels are apparently no problem for the plants under ideal conditions, increased levels are. This makes the *DEAD* motif an important containment mechanism to prevent DRH1 levels from becoming detrimental.

**Figure 6:**
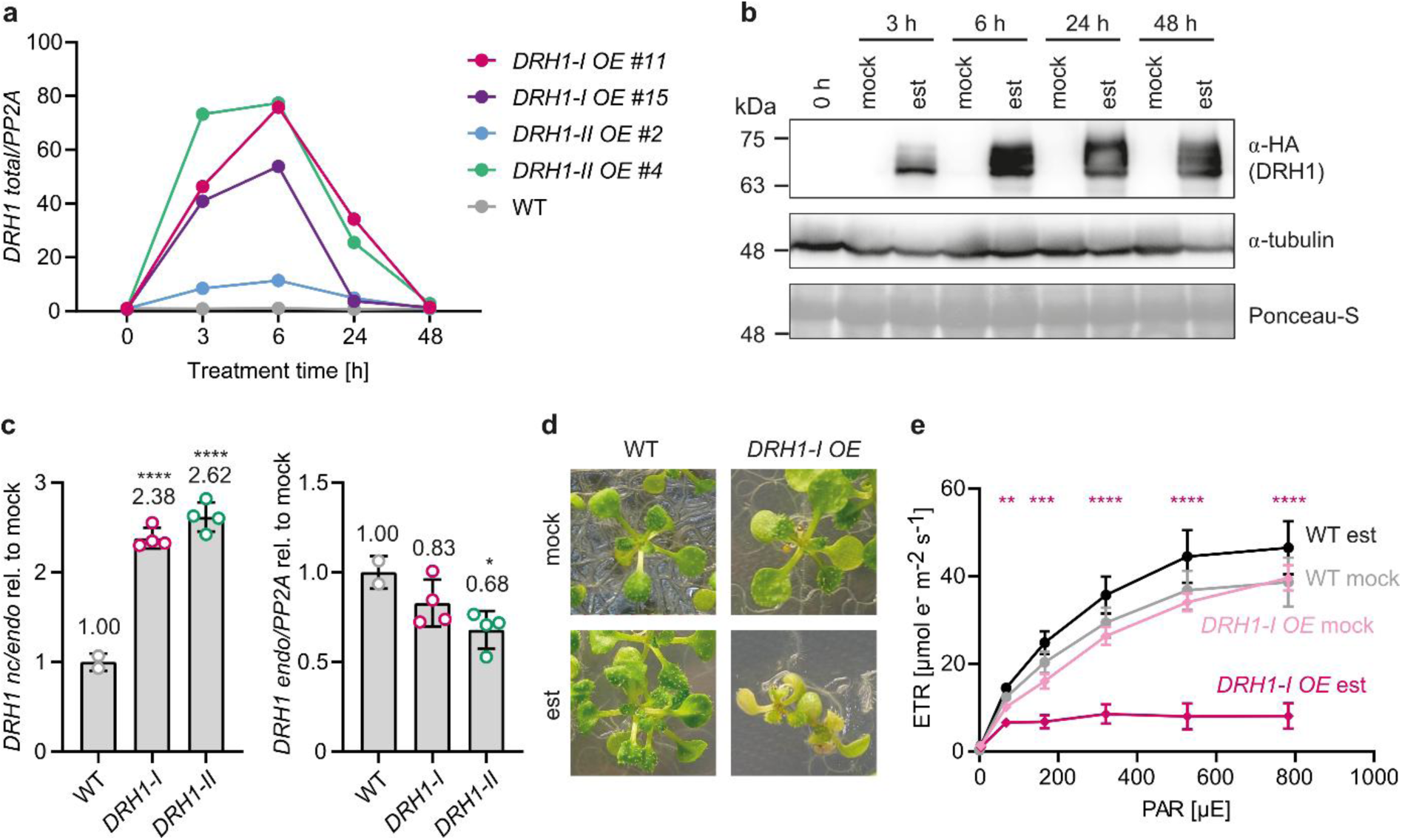
DRH1 overexpression is detrimental for the plant. **a:** Time-course analysis of *DRH1* expression in seedlings of individual *A. thaliana* lines and a WT control after estradiol treatment. Data points represent single biological replicates at a single time point. All values were obtained by qPCR and are relative to the respective mock control. **b:** DRH1 protein expression in seedlings of one representative *A. thaliana* line (*DRH1-I OE #11*) at different time points after mock or estradiol (est) treatment. **c:** Quantitative PCR of endogenous *DRH1* AS ratio and total endogenous transcript levels in WT seedlings and two overexpression lines (*DRH1-I OE* #11, *DRH1-II OE* #4) 6 h after estradiol treatment. All values relative to the respective mock control, the est/mock ratio of transcript levels in WT seedlings was set to 1. *n* = 2 for WT, *n* = 4 for all others. Data points represent biological replicates and are indicated as colored dots. Mean values, standard deviations; asterisks indicate significant change compared to WT (one-way ANOVA followed by Dunnett’s multiple comparisons test, *p < 0.05, ****p < 0.0001). **d:** 14-day-old seedlings (WT or the same *DRH1-I* overexpression line shown in **b-c**) 4 days after mock or est treatment. **e:** Electron transport rate (ETR) in WT seedlings or the same *DRH1-I* overexpression line as in **b-d** 4 days after induction. The same seedlings were measured at multiple light intensities. PAR: photosynthetically active radiation, depicted are mean values (*n* = 5) and standard deviations. Asterisks indicate significant change between est and mock treatment (two-way repeated measures ANOVA followed by Šídák’s multiple comparisons test; **p < 0.01, ***p < 0.001, ****p < 0.0001).

### DRH1 is a major regulator of gene expression

We wondered how circumvention of *DEAD*-mediated control would affect gene expression. Consequently, we performed whole transcriptome sequencing of both mock- and estradiol-treated seedlings carrying the DRH1-I overexpression construct. To capture both early and late responses before the seedlings developed a visible stress phenotype, we analyzed samples 3 h, 6 h and 24 h after DRH1 induction and compared transcriptome patterns between treatments. We found 1,883 DAS and 4,904 DE genes for at least one time point after estradiol treatment (Fig. 7a, SFig. 9a, Supplemental Data S3). The number of events increased with the time of DRH1 overexpression, and the overlap between time points was quite large, indicating that most of the early changes persisted (Fig. 7a). In contrast, the groups of DAS and DE genes showed very little overlap (SFig. 9b), suggesting that the different types of regulation were associated with different target pools. Interestingly, the early response seemed to mainly involve AS, as there were more than twice as many DAS as DE genes 3 h after DRH1 induction (Fig. 7b). However, this changed with time: 6 h after induction, the numbers of significant DAS and DE genes were approximately equal, and after 24 h, DE genes predominated. This suggests that DRH1 has a major impact on AS control, whereas most of the later changes are likely secondary effects that may not be directly caused by DRH1. The direction of regulation also changed with time: 6 h after estradiol treatment, most DE genes were downregulated, whereas after 24 h, the majority showed higher transcript levels than in the mock samples (Fig. 7c).

**Figure 7:**
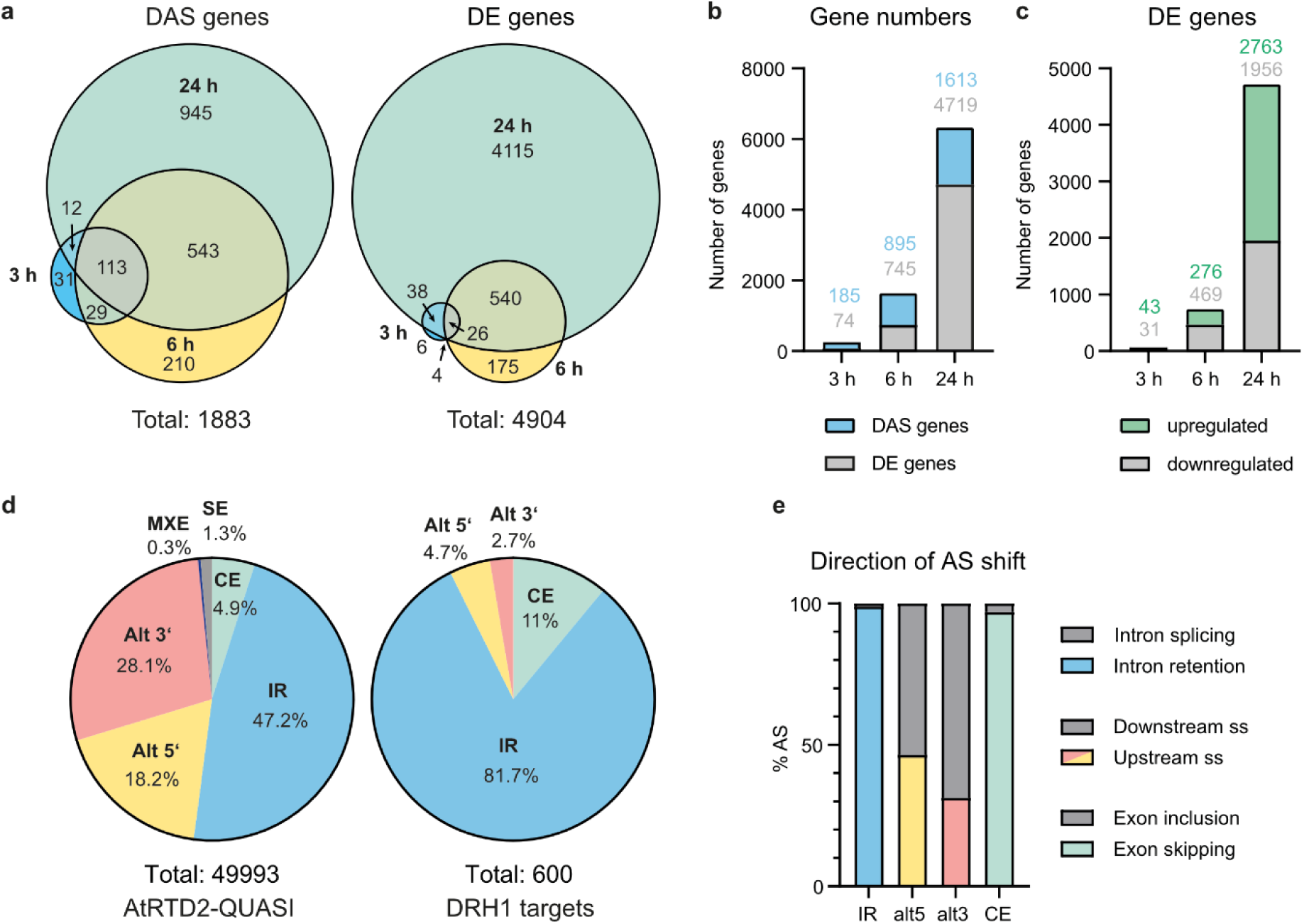
*DRH1* overexpression causes major changes in AS and gene expression. **a:** Venn diagrams showing the numbers and overlaps between significant (p < 0.01) DAS or DE genes obtained for the three time points after induction. Total numbers of significant DAS and DE genes over all time points are given at the bottom. **b:** Numbers of significant DE and DAS genes shown individually for the respective time points. **c:** Numbers of significant DE genes that are up- or downregulated at different time points after *DRH1* induction. **d:** Frequencies of AS event types in the reference transcriptome (AtRTD2-QUASI, left) compared to the events regulated by *DRH1* at the 3 h and 6 h time points (right). Total number of events is given below each chart. MXE: mutually exclusive exons, SE: shifted exon (both alt 5’ and alt 3’ splice site differ). **e:** Direction of AS shift induced by DRH1 overexpression for each AS type.

We tested 10 random DAS genes that already showed a change at the 6 h time point via RT-PCR and could validate all of them (SFig. 9c-d). To find out if DRH1 preferentially regulates a certain type of AS event, we analyzed AS types among DRH1 targets in comparison to the reference transcriptome (AtRTD2-QUASI) ^45^. We compared all reciprocal transcript pairs of a gene (i.e., at least one transcript is upregulated by DRH1 overexpression and at least one decreases) and included all those that differed in a single AS event. DRH1 seems to prefer CE and IR events: They amount to 11% and 81.7% of individual AS events regulated by DRH1, respectively; whereas their frequencies in the reference transcriptome are 4.9% and 47.2% (Fig. 7d). In contrast, alt3 and alt5 splice site events are underrepresented among DRH1 targets. Interestingly, there is a clear directional bias for the regulated CE and IR events: In more than 95% of all cases, DRH1 promotes exon skipping and intron retention, respectively (Fig. 7e).

We next investigated if DRH1 also shows a preference for a specific subset of genes. We performed a Gene Ontology (GO) term enrichment analysis including all significant DAS or DE genes over all three time points (Supplemental Data S4). Among DAS genes, there is an enrichment of RNA-related terms such as “RNA splicing” and “mRNA processing” as well as “nucleus”, “nuclear speck” and “P-body”. In addition, there are terms connected to protein binding/modification, kinase activity, and stress response. DE genes fall into a large number of stress-related categories and include many chloroplast-related terms, suggesting that misregulation of these genes might be responsible for the bleaching observed upon DRH1 induction. Accordingly, DRH1 regulates other RBPs and might also be involved in signaling cascades including stress responses.

Taken together, an excess of DRH1 leads to massive changes in AS and/or gene expression that affect almost a third of the transcriptome, with early responses mostly altering AS and later, possibly secondary effects mainly functioning via differential expression. This establishes DRH1 as a critical regulator of gene expression with implications for various processes in plant development and stress response.

## Discussion

### DRH1 has a major impact on AS and gene expression

Here, we establish the DEAD-box RNA helicase DRH1 as a splicing regulator with more than 1,800 target genes, but also as a major regulator of gene expression (Fig. 7, SFig. 9). This is well in line with the activity of the closest human homologs, DDX5 and DDX17, which were shown to control AS of thousands of genes ^46,47^, but opens the question how an RNA helicase regulates AS and gene expression on such a large scale. An exciting possibility is that DRH1 could alter secondary RNA structures globally and thereby cause the observed changes. A transcriptome-wide *in vivo* structure profiling study in *A. thaliana* found that efficient splicing is associated with single-stranded nucleotides directly at the 5’ splice site and at the branch point, and that base-pairing at the two positions upstream of the 5’ splice site is sufficient to prevent its usage ^9^. This feature was also shown to be involved in alternative splice site choice. Unwinding of dsRNA around alternative 5’ splice sites could thus lead to different AS outcomes. Another possibility is that structural rearrangement of *cis*-acting elements that are involved in AS regulation alters their accessibility for the associated *trans*-acting factors. Such a dependency on RNA structure was shown for several human splicing regulators ^48,49^ and DDX5 can interfere with binding of some RBPs in such a manner ^20,21,46^. It has also been suggested that RNA structure can stand in for other *cis*-regulatory signals. Rapidly evolving exons with a high number of polymorphisms are more likely to be flanked by highly structured introns, possibly compensating for the loss of exonic splicing signals by bringing splice sites into close proximity ^50^. Loss of this supposed safeguarding mechanism due to unregulated helicase activity could therefore lead to severe phenotypes. Furthermore, DRH1 might resolve intermolecular RNA interactions such as miRNAs or snRNAs bound to their targets, which would also lead to large-scale disturbances in gene regulation and AS. For DDX5, it has been proposed that it can unwind the duplex formed between U1 snRNA and a 5’ splice site ^51^, so a similar function of DRH1 would be conceivable. In addition, several DEAD-box helicases were shown to actively remodel RNA-protein complexes by displacing RBPs from their target RNAs, including large and essential assemblies like the exon junction complex or U1 snRNP ^52–54^. Such splicing-inhibitory functions would fit well to the observation that DRH1 mainly promotes IR and exon skipping, i.e., weak splice sites might be less efficiently recognized. Interestingly, another *A. thaliana* DEAD-box helicase, RCF1, was also shown to affect AS of a large number of targets with an overrepresentation of CE events ^55^, suggesting that this could be a general feature of this protein family. Data from DDX5 also indicates that binding might not be completely random. Although the helicase domains mainly interact with the RNA backbone and therefore bind rather unspecifically ^17^, DDX5 was shown to prefer GC-rich sequences mainly at junctions between single- and double-stranded RNA ^47^. Some of these motifs overlapped with binding sites of another splicing regulator, hnRNPA1, suggesting that specific regulation of the helicase could be conferred by other RBPs. Helicase overexpression would then likely oversaturate levels of these RBPs, thus leading to uncontrolled activity. Taken together, global interference with RNA structures and/or RNA-protein complexes by unbalanced activity of DRH1 could explain at least some of the massive changes in gene expression and the resulting stress phenotype that is caused by DRH1 overexpression.

### *DEAD* functions as a sensor for DRH1 helicase activity

The major function of DRH1 in gene expression probably necessitates precise control of its activity. We propose that the structured RNA element *DEAD* within the *DRH1* and *RH46* pre-mRNAs is part of this control system by acting as a sensor for cellular levels of the two helicases. When helicase activity is low, the RNA folds into the predicted hairpin and thus obstructs an alternative splice site located at the base of the stem, leading to almost exclusive splicing towards the coding transcripts (Fig. 8). Translation of these mRNAs results in increased levels of the two helicases in the cell which in turn leads to increased unwinding of the motif, thus facilitating access to the splice site and inducing a splicing shift towards non-coding transcripts. Interestingly, usage of the alternative splice site in *N. benthamiana* requires besides opening of the hairpin also the presence of DRH1 or possibly RH46 (Fig. 4a-b, SFig. 2c, SFig. 3a-b), indicating that the two proteins have a function beyond unwinding such as recruitment of splicing factors or aiding in spliceosome assembly. RH46 can control AS of *DRH1* in *N. benthamiana* (Fig. 4a-b, SFig. 3a-b), but its knockout in *A. thaliana* had no substantial effect on non-coding transcripts of either *DRH1* or *RH46* (SFig. 5e, h). This might be due to generally low expression of *RH46* in seedlings (SFig. 6). In *A. thaliana*, DRH1 would thus negatively regulate itself and its paralog via unwinding of *DEAD*.

**Figure 8:**
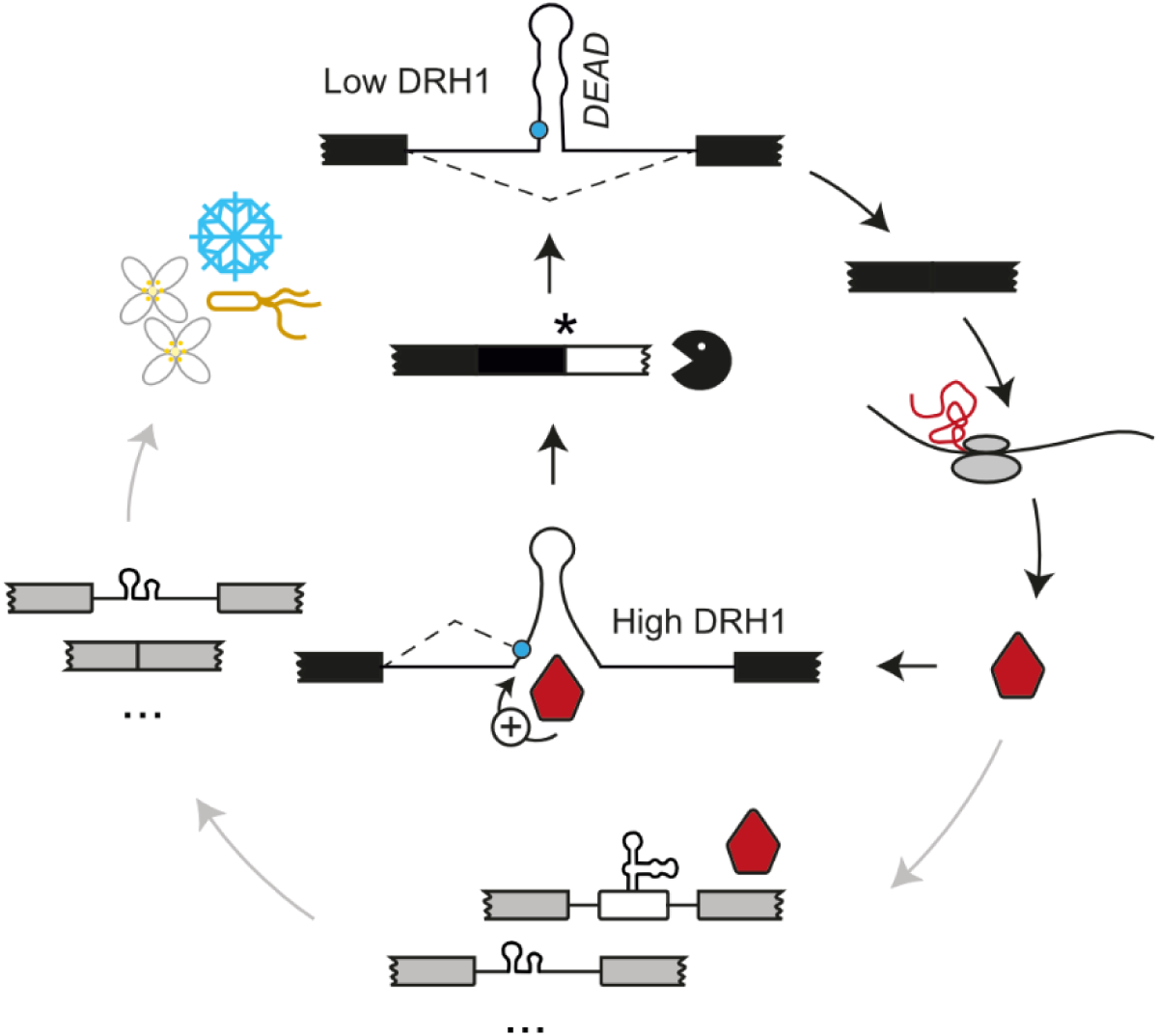
Model of *DEAD*-mediated *DRH1* regulation. Under low DRH1 levels/activity, the *DEAD* motif forms a stem-loop structure, obstructing the alternative 3’ splice site located at the bottom of the stem (blue circle). *DRH1* pre-mRNA is therefore spliced almost exclusively towards the coding variant which is translated into DRH1 protein. The increase in DRH1 levels leads to unwinding of the *DEAD* motif, facilitating access to the splice site and splicing towards the non-coding variants. This negative feedback loop balances DRH1 expression (black arrows). Moreover, the helicase can regulate AS and gene expression of multiple downstream targets, mainly repressing splice site usage at IR and CE events (grey arrows). This is possibly accomplished via global rearrangement of RNA secondary structure and affects various processes, including flowering time and expression of stress-related genes.

DEAD-box RNA helicases are reported to efficiently unwind a maximum duplex length of 10-15 bp ^56^. The *DEAD* motif with its 20 bp long stem is outside of this range. However, opening of only the bottom half of the *DEAD* stem dramatically increased usage of the 3’ splice site (Fig. 3), suggesting that this is either sufficient to allow spliceosome access, or it destabilizes the remaining half of the duplex. At least some DEAD-box helicases can start the unwinding reaction from a double-stranded region ^18^, meaning that DRH1 could also locally disrupt base-pairing around the splice site without affecting the ends of the helix. However, the length of *DEAD* is very conserved (Fig. 1b), so it is likely that a longer duplex is needed for proper function. For example, the higher number of base-pairing nucleotides might be needed to stabilize the structure in its natural pre-mRNA context, preventing it from dissociating without activity of a helicase.

Interestingly, *DEAD* seems to act as a sensor for activity of DRH1 and RH46 only, as closely related DEAD-box helicases did not or only weakly promote usage of the alternative 3’ splice site within *DRH1* (Fig. 4). Either the other proteins do not unwind *DEAD*, or they lack the ability to induce alternative splice site choice. This specificity is at least partially conferred by the C-terminal RGG/RG motif of DRH1, a domain type that is typically disordered and implicated in various functions, including RNA and protein binding, nuclear localization, translation regulation, and phase separation and speckle formation ^57^. The C-termini of the two paralogs could therefore mediate specific recognition of *DEAD* or interaction with unknown factors that enable them to influence splice site choice. On the other hand, they might also confer efficient localization to the nucleus or mRNP granules that facilitate splicing. Still, the question remains why out of the 56 DEAD-box RNA helicases in *A. thaliana* only these two are regulated by *DEAD*. Overexpression of human DDX5 is associated with cancer ^31^, whereas in plants, elevated levels of several other RNA helicases were not detrimental, but on the contrary mostly conferred enhanced stress tolerance through diverse mechanisms ^58^. Only one member of the *A. thaliana* DEAD-box family, RCF1/SHI2, has been implicated in AS and gene expression on a similar scale as DRH1 ^55,59^, however, overexpression of this helicase has not been tested. It is therefore possible that a large impact on the transcriptome is not common to many helicases and necessitates a safeguarding mechanism specifically for DRH1 and RH46. The evolution of *DEAD* as such a mechanism that employs both AS and structural rearrangement would then present an ingenious way to specifically balance DRH1 and RH46 with minimal cost.

The results presented here suggest that a balanced structural RNA environment is critical for cellular function. The activity of factors that can alter secondary structure on a larger scale must thus be tightly controlled to avoid misregulation of gene expression. Helicase-sensing and -regulating elements such as *DEAD* might therefore be more common in eukaryotic transcriptomes, acting as safeguards and modulators of RNA processing.

## Materials and Methods

### Plant cultivation and transformation

For growing *Arabidopsis thaliana* Col-0 in sterile culture, seeds were surface-sterilized with 3.75% NaOCl and 0.01% Triton X-100, stratified at 4 °C for 1-2 days and germinated on half-strength Murashige and Skoog medium including vitamins (Duchefa) containing 2% sucrose and 0.8% plant agar. For segregating mutant lines, 25 µg/mL Kanamycin or 5 µg/mL Basta Finale (Bayer AG) were added to the medium. For propagation, seedlings were transferred to soil and further cultivated in a climate chamber. All *A. thaliana* plants were kept under long-day conditions (16 h light/22 °C, 8 h dark/20 °C) at ∼100 µE white light and 60% relative humidity. *Nicotiana benthamiana* was cultivated on soil in a climate chamber (16 h light/24 °C, 8 h dark/22 °C, ∼120 µE white light, 60% relative humidity).

*A. thaliana* was transformed via the floral dip method as previously described ^60^. Mutants were selected based on their resistance either on half-strength Murashige and Skoog medium (including vitamins) containing 0.8% plant agar, 50 µg/mL Kanamycin and 200 µg/mL Cefotaxime or by spraying two-week-old soil-grown seedlings with 40 µg/mL Basta Finale (Bayer AG). Transformants were transferred to soil and cultivated until seed set. CRISPR lines were selected via the fluorescence-accumulating seed technology (FAST) method ^61^ under a Leica M205FCA fluorescence stereomicroscope (excitation: 541-551 nm, emission: 565-605 nm) and directly germinated on soil. Later generations were tested for fluorescence to select for transgene-free plants. *N. benthamiana* was transformed via a previously described leaf infiltration assay ^15^ with the following modifications: mOrange2 ^62^ was used as normalization fluorophore, and leaf material was harvested two days after infiltration for RNA isolation and three days after infiltration for protein extraction.

Crossing of the *rh14* line (SALK_073018, obtained from NASC) with a homozygous *rh46* CRISPR line was performed by emasculating young buds and pollinating them on the next day with pollen from the crossing partner.

### Determination of flowering time

Seeds were stratified at 4 °C for 3 days, then sown onto soil and incubated under the conditions specified above. Flowering time was defined as the day after sowing on which the inflorescence reached 1 cm in length.

### Genotyping of plant mutants

The genetic status of *rh14*, *rh46* and *rh14rh46* plants was confirmed by isolation of genomic DNA and genotyping PCR. ∼100 mg ground plant tissue was suspended in 500 µL buffer (200 mM Tris-HCl pH 9, 400 mM LiCl, 25 mM EDTA, 1% SDS) and centrifuged for 5 min at 15,000 x g. The same volume of isopropanol was added to the supernatant, incubated for 5 min at room temperature and gDNA precipitated by centrifugation for 10 min at 15,000 x g. Pellet was washed with 70% ethanol, dried and resuspended in ½ TE buffer (5 mM Tris-HCl pH 8, 0.5 mM EDTA). Genotyping PCR was carried out according to standard procedures with the primers indicated in Supplemental Data S5. For CRISPR lines, desired mutations and predicted off-targets were analyzed by purification of PCR fragments (GeneJET PCR Purification Kit, Thermo Fisher Scientific) and Sanger sequencing. To identify the stop codon in the *DRH1* transcript generated in *rh14* plants, 3’ RACE (Rapid Amplification of cDNA Ends) was performed with GeneRacer™ Kit (Invitrogen) using the nested PCR protocol with the following modifications: SuperScript II (Invitrogen) was used for reverse transcription, and 3’ RACE PCR products were cloned into pGEM-T (Promega) for sequencing.

### Estradiol treatment

DRH1 overexpression was induced by spraying 10-day-old seedlings grown in sterile culture with deionized water containing either 50 µM β-estradiol (Merck, dissolved in DMSO) or the corresponding amount of solvent. 0.02% Silwet L-77 was added to the spraying solution for better tissue penetration. Plates were re-sealed and kept in the climate chamber for up to 4 days.

### ETR measurement

Electron transport rate (ETR) was measured in a Hexagon-Imaging-PAM chlorophyll fluorometer (Heinz Walz GmbH). Dark-acclimated seedlings were subjected to six actinic light pulses with increasing intensities (4, 68, 166, 321, 527 and 783 µE) in 20 sec intervals. Measuring light intensity and frequency were set to 1 (lowest setting). ETR was calculated with the following formula: ETR = Y(II)*PAR*0.5*0.84. Y(II) is the effective photosystem II quantum yield and calculated as Y(II) = (Fm’ – F)/Fm’, with Fm’ being the maximum fluorescence yield at saturating light conditions and F representing the steady-state fluorescence yield (averaged over 3 seconds after a light pulse). ETR values were averaged over 5 areas of interest per line and treatment, each encompassing 1-3 seedlings.

### Cloning procedures

An overview of all constructs used in this study can be found in Supplemental Data S6. All reporter constructs were in the pBinAR vector background ^63^ and were created using the previously published *TFIIIA* reporter ^64^ as template. The 5’ sequence of the gene of interest was amplified from *A. thaliana* genomic DNA using the primers indicated in Supplemental Data S5 and inserted into the vector in place of the *TFIIIA* sequence by *Kpn*I/*Xba*I digest and T4 ligation. The CDS constructs of *UPF1* and *mOrange2* were previously described ^65,66^. The ones of *DRH1-I*, *DRH1-II*, *RH46-I*, *RH46-II*, *RH30*, and *RH20* used for leaf infiltration were also in the pBinAR vector background and were cloned by PCR amplification from cDNA and ligation via *Kpn*I/*Xba*I (*DRH1*) or *Xba*I/*Sal*I (all others). Note that the constructs for *RH46-I*, *RH46-II*, *RH30,* and *RH20* exist with and without an N-terminal single HA tag, the tagged ones were only used for the data in Fig. 4d and f and SFig. 3c. *RH20* and *RH30* constructs with the C-terminal RGG domain of *DRH1* were created via overlap PCR, subcloned into pCR-Blunt-II-TOPO (Invitrogen) and cloned into pBinAR via *Xba*I/*Sal*I. To generate the CDS constructs of DRH1 helicase domain mutants, a version of pBinAR carrying an N-terminal triple HA tag with start codon was generated by cloning of an oligo annealing product into the vector via *Kpn*I/*Bam*HI digest. *DRH1* inserts were then cloned into this plasmid by digest with *Bam*HI/*Xba*I. Mutations of the *DEAD* motif and the *DRH1* helicase domain were introduced by subcloning the respective construct into pGEM-T (Promega) and performing PCR mutagenesis with the primers indicated in Supplemental Data S5.

*DRH1* overexpression and genomic complementation constructs were created via the GreenGate system using several pre-existing modules as listed in Supplemental Data S6 ^67,68^. The following modules were cloned by PCR amplification from *A. thaliana* cDNA and *Bsa*I-mediated ligation into pGGC000: *DRH1-I CDS* (CRB01), *DRH1-II CDS* (CRB02). The N-terminal 3xHA tag module was created via oligo annealing and ligation into pGEM-T (Promega). For *pUBQ:DRH1* genomic constructs, the genomic DNA of *DRH1* was divided into three parts: 5’ region with single HA tag integrated before the start codon (C1), region from start codon to exon 6 (C2), and 3’ region (C3). Mutations of *DEAD* were introduced into C2 modules by using the respective reporter constructs as templates. Fragments were cloned into pGEM-T (Promega) and combined to form one CDS module. The exact configuration of all GreenGate constructs can be found in Supplemental Data S6. In all *DRH1* sequences used for GreenGate cloning, the two internal *Bsa*I sites were removed by PCR mutagenesis with the primers indicated in Supplemental Data S5.

The CRISPR-Cas9 construct to create *rh46* knockout plants was generated using a published GoldenGate cloning system ^69^. The sgRNAs indicated in Supplemental Data S5 were designed using CRISPR-P 2.0 ^70^ and cloned into the pDGE652 vector under control of the *A. thaliana* U6-26 promoter fragment as described by Stuttmann et al. (2021). Note that the final construct includes an sgRNA against *DRH1* which turned out to be nonfunctional as no corresponding mutants could be identified.

For identification of splicing variants, the PCR fragments were cloned into pGEM-T (Promega) and sequenced via Sanger sequencing.

### RNA isolation and RT-PCR

RNA was isolated from ∼100 mg ground plant tissue using the EURx Universal RNA Purification Kit (Roboklon), including an on-column DNase I digest as specified in the manufacturer’s instructions. Reverse transcription was performed with AMV Reverse Transcriptase Native (Roboklon) or SuperScript II (Invitrogen) using a dT_20_ primer. RT-PCR was carried out according to standard procedures, PCR products were separated via agarose gel electrophoresis and visualized by ethidium bromide staining.

### Transcript quantification

RT-PCR products were quantified on an Agilent Bioanalyzer 2100 using the DNA1000 protocol. Quantification of transcripts in *DRH1* misexpression lines took place via quantitative PCR using MESA-BLUE qPCR Mastermix Plus for SYBR Assay, no ROX (Eurogentec) in a CFX384 cycler (BioRad). Primers are indicated in Supplemental Data S5. All reactions were performed in triplicate, and a melt curve was added. Data was analyzed using the relative standard curve method, a cDNA dilution series was included to measure primer efficiencies. Transcript levels were normalized to the housekeeping gene *AT1G13320* (*PP2A* catalytic subunit) or to *GFP* (reporter construct).

### Whole-transcriptome sequencing

150 ng total RNA of 2 biological replicates per sample were used. RNA quality was assessed by RNA integrity number (RIN) measurement on an Agilent Bioanalyzer 2100 using the RNA6000 Nano protocol. Concentration was measured on a Qubit 4 fluorometer (Invitrogen) using the RNA broad range assay. Poly-A selection as well as generation and sequencing of strand-specific cDNA libraries took place at Eurofins Genomics on a NovaSeq 6000 S4 system, generating a minimum of 25 million read pairs (2 x 150 nt) per sample. Reads were trimmed with BBDuk (Bushnell B., www.sourceforge.net/projects/bbmap) and mapped to the reference transcriptome AtRTD2-QUASI ^45^ using Salmon ^71^. Data was analyzed using the 3D RNA-seq App ^72^ with the following cut-off values: p > 0.01, log2 fold change (DE) > 1, deltaPS (DAS) > 0.1; lowly expressed transcripts were filtered using a cut-off value of 4 counts per million (cpm) in at least 1 sample. Gene Ontology enrichment analysis was performed with DAVID ^73^.

### Event type analysis

Event types were determined using a custom script (available at https://github.com/MaxiSack/ReciprocalEventIdentification). All pairs of transcripts (for each regulated gene) that are reciprocally regulated by DRH1 (i.e., one transcript is upregulated by DRH1 overexpression and one decreases) for the 3 h and 6 h time points were identified. Using the intron-exon annotation provided by the reference transcriptome (AtRTD2-QUASI), we inferred which AS event connects the two. For the sake of simplicity and to avoid ambiguity, we only investigated those pairs of transcripts that are connected through exactly one AS event. We considered the following events and inferred them as such:

1. If all exons were identical, except that one transcript had an additional exon, it was counted as cassette exon and the transcripts are therefore connected through an exon skipping/inclusion event.
2. If there were two exons in one transcript and one in the other that are not present in the other respective transcript, and if then the two exons were next to another and their outer borders matched the borders of the one exon in the other transcript, it was counted as an intron retention/splicing event.
3. If there was one exon in each transcript that shared either their 5’ or 3’ border, but not the other, it was counted as an alternative 3’ or 5’ splice site event, respectively.

Any pair of reciprocally regulated transcripts whose exons differed by more than one of these three cases was ignored. Finally, it was noted in which direction the AS event happened in response to DRH1.

### Protein extraction and fluorescence assay

Proteins were isolated from ∼100 mg ground tissue by suspension in 300 µL buffer (50 mM Tris pH 7.5, 150 mM NaCl, 0.1% Tween, 0.1% β-mercaptoethanol) and centrifugation at 4 °C and 15,000 x g for 15 min. Supernatant was used for Western blot or fluorescence measurement. For the latter, 100 µL of supernatant was transferred to a 96-well plate (Greiner, black, flat bottom) and fluorescence of EGFP and mOrange2 measured in a Tecan M1000 plate reader with the following settings: exc. 485 ± 7 nm, em. 520 ± 5 nm (EGFP), exc. 520 ± 5 nm, em. 600 ± 5 nm (mOrange2). mOrange2 fluorescence was used to correct for variances in transformation efficiency.

### Western Blot

10-15 µg of protein were denatured by addition of 10x SDS buffer (300 mM Tris pH 6.8, 50% (w/v) glycerol, 5% (w/v) SDS, 15 µM bromophenol blue, 100 mM DTT) and incubation at 95 °C for 5 min. Proteins were separated on a 10 % SDS-PAGE gel using a Tris-glycine buffer (25 mM Tris, 200 mM glycine, 0.1% SDS) and transferred to a nitrocellulose membrane in a similar buffer (25 mM Tris, 140 mM glycine, 20% EtOH). Membrane was washed in TBS-T (10 mM Tris pH 7.5, 150 mM NaCl, 0.05% Tween-20), blocked for 1 h at room temperature in TBS-T + 5% skimmed milk and washed again before application of antibody (1 h at room temperature or overnight at 4 °C). The following primary antibodies were used in this study: α-HA (rat, Roche 11867423001), α-tubulin alpha (rabbit, Agrisera AS10 680), α-FLAG (mouse, Merck F3165). The following secondary antibodies were used: α-rat-PRX (goat, Thermo Fisher Scientific 31470), α-rabbit-PRX (goat, Merck A6154), α-mouse-PRX (goat, Merck A9917). Detection was performed with Western Blot Hyper HRP substrate (TaKaRa) in a Fusion FX imager (Vilber Lourmat).

### Statistics

All statistics except those implemented in 3D RNA-seq ^72^ were calculated with GraphPad Prism 9.4.0 and are described in the figure legends and in Supplemental Data S7. Data sets were tested for normal distribution using Shapiro-Wilk test and for equal variance using Brown-Forsythe test. For comparison of more than 2 populations, one-way ANOVA followed by Dunnett’s multiple comparisons test was used in case of normality and Kruskal-Wallis test followed by Dunn’s multiple comparisons test if normality criterion was not met. For normally distributed data with unequal variance, Brown-Forsythe and Welch ANOVA followed by Dunnett’s T3 multiple comparisons test was used. Comparison of ETR curves (which were generated by repeated measurements of the same samples) were performed using two-way repeated measures ANOVA followed by Šídák’s multiple comparisons test. In cases where data was normalized to an internal control that therefore lacked variance, one-sample t-test against a hypothetical mean of 1 was performed.

## Code availability

Custom code used for analysis of AS event types is available on GitHub (https://github.com/MaxiSack/ReciprocalEventIdentification).

## Acknowledgements

We are grateful to Marc Gebauer for technical assistance.

## Author contributions

Conceptualization: AW and RB; Investigation: RB, JB, MR, NR; Formal Analysis: CE, SLH, MS, ZW; Writing: RB and AW; Supervision: AW and ZW; Funding acquisition: AW and ZW.

## Supplemental material

- Supplemental Figures
- Supplemental File 1: *DEAD* alignment
- Supplemental Data S1: List of *DEAD* homologs found
- Supplemental Data S2: RNA-seq results of the knockout lines
- Supplemental Data S3: RNA-seq results of *DRH1* overexpression lines
- Supplemental Data S4: GO term analysis
- Supplemental Data S5: List of DNA oligos
- Supplemental Data S6: Overview of cloned constructs
- Supplemental Data S7: Statistics

## References

1. Chung, B. Y. W. et al. An RNA thermoswitch regulates daytime growth in Arabidopsis. Nat Plants 6, 522–532; 10.1038/s41477-020-0633-3 (2020).

2. Gosai, S. J. et al. Global analysis of the RNA-protein interaction and RNA secondary structure landscapes of the Arabidopsis nucleus. Mol Cell 57, 376–388; 10.1016/j.molcel.2014.12.004 (2015).

3. Saldi, T., Fong, N. & Bentley, D. L. Transcription elongation rate affects nascent histone pre-mRNA folding and 3’ end processing. Genes Dev 32, 297–308; 10.1101/gad.310896.117 (2018).

4. Steinert, H. et al. Pausing guides RNA folding to populate transiently stable RNA structures for riboswitch-based transcription regulation. eLife 6; 10.7554/eLife.21297 (2017).

5. Sun, L. et al. RNA structure maps across mammalian cellular compartments. Nat Struct Mol Biol 26, 322–330; 10.1038/s41594-019-0200-7 (2019).

6. Wang, X.-W., Liu, C.-X., Chen, L.-L. & Zhang, Q. C. RNA structure probing uncovers RNA structure-dependent biological functions. Nat Chem Biol 17, 755–766; 10.1038/s41589-021-00805-7 (2021).

7. Taylor, K. & Sobczak, K. Intrinsic Regulatory Role of RNA Structural Arrangement in Alternative Splicing Control. Int J Mol Sci 21, 5161; 10.3390/ijms21145161 (2020).

8. Cao, X., Zhang, Y., Ding, Y. & Wan, Y. Identification of RNA structures and their roles in RNA functions. Nat Rev Mol Cell Biol 25, 784–801; 10.1038/s41580-024-00748-6 (2024).

9. Liu, Z. et al. In vivo nuclear RNA structurome reveals RNA-structure regulation of mRNA processing in plants. Genome Biol 22, 11; 10.1186/s13059-020-02236-4 (2021).

10. Wan, Y. et al. Landscape and variation of RNA secondary structure across the human transcriptome. Nature 505, 706–709; 10.1038/nature12946 (2014).

11. Shepard, P. J. & Hertel, K. J. Conserved RNA secondary structures promote alternative splicing. RNA 14, 1463–1469; 10.1261/rna.1069408 (2008).

12. Xu, B., Shi, Y., Wu, Y., Meng, Y. & Jin, Y. Role of RNA secondary structures in regulating Dscam alternative splicing. Biochimica et biophysica acta. Gene regulatory mechanisms 1862, 194381; 10.1016/j.bbagrm.2019.04.008 (2019).

13. Bocobza, S. et al. Riboswitch-dependent gene regulation and its evolution in the plant kingdom. Genes Dev 21, 2874–2879; 10.1101/gad.443907 (2007).

14. Cheah, M. T., Wachter, A., Sudarsan, N. & Breaker, R. R. Control of alternative RNA splicing and gene expression by eukaryotic riboswitches. Nature 447, 497–500; 10.1038/nature05769 (2007).

15. Wachter, A. et al. Riboswitch control of gene expression in plants by splicing and alternative 3’ end processing of mRNAs. Plant Cell 19, 3437–3450; 10.1105/tpc.107.053645 (2007).

16. Jones, A. N. et al. Modulation of pre-mRNA structure by hnRNP proteins regulates alternative splicing of MALT1. Sci Adv 8, eabp9153; 10.1126/sciadv.abp9153 (2022).

17. Fairman-Williams, M. E., Guenther, U.-P. & Jankowsky, E. SF1 and SF2 helicases: family matters. Curr Opin Struct Biol 20, 313–324; 10.1016/j.sbi.2010.03.011 (2010).

18. Yang, Q., Del Campo, M., Lambowitz, A. M. & Jankowsky, E. DEAD-box proteins unwind duplexes by local strand separation. Mol Cell 28, 253–263; 10.1016/j.molcel.2007.08.016 (2007).

19. Linder, P. & Jankowsky, E. From unwinding to clamping - the DEAD box RNA helicase family. Nat Rev Mol Cell Biol 12, 505–516; 10.1038/nrm3154 (2011).

20. Laurent, F.-X. et al. New function for the RNA helicase p68/DDX5 as a modifier of MBNL1 activity on expanded CUG repeats. Nucleic acids research 40, 3159–3171; 10.1093/nar/gkr1228 (2012).

21. Camats, M., Guil, S., Kokolo, M. & Bach-Elias, M. P68 RNA helicase (DDX5) alters activity of cis- and trans-acting factors of the alternative splicing of H-Ras. PLOS ONE 3, e2926; 10.1371/journal.pone.0002926 (2008).

22. Kar, A. et al. RNA helicase p68 (DDX5) regulates tau exon 10 splicing by modulating a stem-loop structure at the 5’ splice site. Mol Cell Biol 31, 1812–1821; 10.1128/MCB.01149-10 (2011).

23. Yeh, F.-L. et al. Activation of Prp28 ATPase by phosphorylated Npl3 at a critical step of spliceosome remodeling. Nat Commun 12, 3082; 10.1038/s41467-021-23459-4 (2021).

24. Taylor, K. et al. Modulatory role of RNA helicases in MBNL-dependent alternative splicing regulation. Cell. Mol. Life Sci. 80, 335; 10.1007/s00018-023-04927-0 (2023).

25. Li, X. et al. Functions and mechanisms of RNA helicases in plants. J Exp Bot 74, 2295–2310; 10.1093/jxb/erac462 (2023).

26. Sack, M. et al. Identification and characterization of new structured RNA classes in plants. RNA Biology 22, 1–16; 10.1080/15476286.2025.2523696 (2025).

27. Burgess, D. & Freeling, M. The most deeply conserved noncoding sequences in plants serve similar functions to those in vertebrates despite large differences in evolutionary rates. Plant Cell 26, 946–961; 10.1105/tpc.113.121905 (2014).

28. O’Leary, N. A. et al. Reference sequence (RefSeq) database at NCBI: current status, taxonomic expansion, and functional annotation. Nucleic Acids Res 44, D733–45; 10.1093/nar/gkv1189 (2016).

29. Müller-McNicoll, M., Rossbach, O., Hui, J. & Medenbach, J. Auto-regulatory feedback by RNA-binding proteins. J Mol Cell Biol 11, 930–939; 10.1093/jmcb/mjz043 (2019).

30. Ohtani, M. & Wachter, A. NMD-Based Gene Regulation-A Strategy for Fitness Enhancement in Plants? Plant Cell Physiol 60, 1953–1960; 10.1093/pcp/pcz090 (2019).

31. Xing, Z., Ma, W. K. & Tran, E. J. The DDX5/Dbp2 subfamily of DEAD-box RNA helicases. Wiley Interdiscip Rev RNA 10, e1519; 10.1002/wrna.1519 (2019).

32. Jalal, C., Uhlmann-Schiffler, H. & Stahl, H. Redundant role of DEAD box proteins p68 (Ddx5) and p72/p82 (Ddx17) in ribosome biogenesis and cell proliferation. Nucleic Acids Res 35, 3590–3601; 10.1093/nar/gkm058 (2007).

33. Zhang, H. et al. RNA helicase DEAD box protein 5 regulates Polycomb repressive complex 2/Hox transcript antisense intergenic RNA function in hepatitis B virus infection and hepatocarcinogenesis. Hepatology 64, 1033–1048; 10.1002/hep.28698 (2016).

34. Chicois, C. et al. The UPF1 interactome reveals interaction networks between RNA degradation and translation repression factors in Arabidopsis. Plant J 96, 119–132; 10.1111/tpj.14022 (2018).

35. Palm, D. et al. Plant-specific ribosome biogenesis factors in Arabidopsis thaliana with essential function in rRNA processing. Nucleic Acids Res 47, 1880–1895; 10.1093/nar/gky1261 (2019).

36. Sulkowska, A. et al. RNA Helicases from the DEA(D/H)-Box Family Contribute to Plant NMD Efficiency. Plant Cell Physiol 61, 144–157; 10.1093/pcp/pcz186 (2020).

37. Okanami, M., Meshi, T. & Iwabuchi, M. Characterization of a DEAD box ATPase/RNA helicase protein of Arabidopsis thaliana. Nucleic Acids Res 26, 2638–2643; 10.1093/nar/26.11.2638 (1998).

38. Klepikova, A. V., Kasianov, A. S., Gerasimov, E. S., Logacheva, M. D. & Penin, A. A. A high resolution map of the Arabidopsis thaliana developmental transcriptome based on RNA-seq profiling. Plant J 88, 1058–1070; 10.1111/tpj.13312 (2016).

39. Nakabayashi, K., Okamoto, M., Koshiba, T., Kamiya, Y. & Nambara, E. Genome-wide profiling of stored mRNA in Arabidopsis thaliana seed germination: epigenetic and genetic regulation of transcription in seed. Plant J 41, 697–709; 10.1111/j.1365-313X.2005.02337.x (2005).

40. Schmid, M. et al. A gene expression map of Arabidopsis thaliana development. Nat Genet 37, 501–506; 10.1038/ng1543 (2005).

41. Zuo, J., Niu, Q. W. & Chua, N. H. Technical advance: An estrogen receptor-based transactivator XVE mediates highly inducible gene expression in transgenic plants. Plant J 24, 265–273; 10.1046/j.1365-313x.2000.00868.x (2000).

42. Dias, M. C. & Brüggemann, W. Limitations of photosynthesis in Phaseolus vulgaris under drought stress: gas exchange, chlorophyll fluorescence and Calvin cycle enzymes. Photosynthetica 48, 96–102; 10.1007/s11099-010-0013-8 (2010).

43. Pan, C., Ahammed, G. J., Li, X. & Shi, K. Elevated CO2 Improves Photosynthesis Under High Temperature by Attenuating the Functional Limitations to Energy Fluxes, Electron Transport and Redox Homeostasis in Tomato Leaves. Front Plant Sci 9, 1739; 10.3389/fpls.2018.01739 (2018).

44. Wong, S.-L., Chen, C.-W., Huang, M.-Y. & Weng, J.-H. Relationship between photosynthetic CO2 uptake rate and electron transport rate in two C4 perennial grasses under different nitrogen fertilization, light and temperature conditions. Acta Physiol Plant 36, 849–857; 10.1007/s11738-013-1463-y (2014).

45. Zhang, R. et al. A high quality Arabidopsis transcriptome for accurate transcript-level analysis of alternative splicing. Nucleic acids research 45, 5061–5073; 10.1093/nar/gkx267 (2017).

46. Dardenne, E. et al. RNA helicases DDX5 and DDX17 dynamically orchestrate transcription, miRNA, and splicing programs in cell differentiation. Cell Reports 7, 1900–1913; 10.1016/j.celrep.2014.05.010 (2014).

47. Lee, Y. J., Wang, Q. & Rio, D. C. Coordinate regulation of alternative pre-mRNA splicing events by the human RNA chaperone proteins hnRNPA1 and DDX5. Genes Dev 32, 1060–1074; 10.1101/gad.316034.118 (2018).

48. Jolma, A. et al. Binding specificities of human RNA-binding proteins toward structured and linear RNA sequences. Genome Res 30, 962–973; 10.1101/gr.258848.119 (2020).

49. Dominguez, D. et al. Sequence, Structure, and Context Preferences of Human RNA Binding Proteins. Mol Cell 70, 854–867.e9; 10.1016/j.molcel.2018.05.001 (2018).

50. Lin, C.-L., Taggart, A. J. & Fairbrother, W. G. RNA structure in splicing: An evolutionary perspective. RNA Biol 13, 766–771; 10.1080/15476286.2016.1208893 (2016).

51. Lin, C., Yang, L., Yang, J. J., Huang, Y. & Liu, Z.-R. ATPase/helicase activities of p68 RNA helicase are required for pre-mRNA splicing but not for assembly of the spliceosome. Mol Cell Biol 25, 7484–7493; 10.1128/MCB.25.17.7484-7493.2005 (2005).

52. Bowers, H. A. et al. Discriminatory RNP remodeling by the DEAD-box protein DED1. RNA 12, 903–912; 10.1261/rna.2323406 (2006).

53. Fairman, M. E. et al. Protein displacement by DExH/D “RNA helicases” without duplex unwinding. Science 304, 730–734; 10.1126/science.1095596 (2004).

54. Lund, M. K. & Guthrie, C. The DEAD-box protein Dbp5p is required to dissociate Mex67p from exported mRNPs at the nuclear rim. Mol Cell 20, 645–651; 10.1016/j.molcel.2005.10.005 (2005).

55. Xu, C. et al. The DEAD-box helicase RCF1 plays roles in miRNA biogenesis and RNA splicing in Arabidopsis. Plant J 116, 144–160; 10.1111/tpj.16366 (2023).

56. Jarmoskaite, I. & Russell, R. DEAD-box proteins as RNA helicases and chaperones. Wiley Interdiscip Rev RNA 2, 135–152; 10.1002/wrna.50 (2011).

57. Chowdhury, M. N. & Jin, H. The RGG motif proteins: Interactions, functions, and regulations. Wiley Interdiscip Rev RNA 14, e1748; 10.1002/wrna.1748 (2023).

58. Nidumukkala, S., Tayi, L., Chittela, R. K., Vudem, D. R. & Khareedu, V. R. DEAD box helicases as promising molecular tools for engineering abiotic stress tolerance in plants. Critical Reviews in Biotechnology 39, 395–407; 10.1080/07388551.2019.1566204 (2019).

59. Wang, B. et al. The DEAD-box RNA helicase SHI2 functions in repression of salt-inducible genes and regulation of cold-inducible gene splicing. J Exp Bot 71, 1598–1613; 10.1093/jxb/erz523 (2020).

60. Clough, S. J. & Bent, A. F. Floral dip: a simplified method for Agrobacterium-mediated transformation of Arabidopsis thaliana. Plant J 16, 735–743; 10.1046/j.1365-313x.1998.00343.x (1998).

61. Shimada, T. L., Shimada, T. & Hara-Nishimura, I. A rapid and non-destructive screenable marker, FAST, for identifying transformed seeds of Arabidopsis thaliana. Plant J 61, 519–528; 10.1111/j.1365-313X.2009.04060.x (2010).

62. Shaner, N. C. et al. Improving the photostability of bright monomeric orange and red fluorescent proteins. Nat Methods 5, 545–551; 10.1038/nmeth.1209 (2008).

63. Höfgen, R. & Willmitzer, L. Biochemical and genetic analysis of different patatin isoforms expressed in various organs of potato (Solanum tuberosum). Plant Sci 66, 221–230; 10.1016/0168-9452(90)90207-5 (1990).

64. Hammond, M. C., Wachter, A. & Breaker, R. R. A plant 5S ribosomal RNA mimic regulates alternative splicing of transcription factor IIIA pre-mRNAs. Nat Struct Mol Biol 16, 541–549; 10.1038/nsmb.1588 (2009).

65. Kesarwani, A. K. et al. Multifactorial and Species-Specific Feedback Regulation of the RNA Surveillance Pathway Nonsense-Mediated Decay in Plants. Plant Cell Physiol 60, 1986–1999; 10.1093/pcp/pcz141 (2019).

66. Reinhardt, M. et al. The structured mRNA element 45ABC mediates auto- and cross-regulation of RBP45 genes via alternative splicing. bioRxiv, 2025.11.25.690383; 10.1101/2025.11.25.690383 (2025).

67. Lampropoulos, A. et al. GreenGate---a novel, versatile, and efficient cloning system for plant transgenesis. PLOS ONE 8, e83043; 10.1371/journal.pone.0083043 (2013).

68. Saile, J. et al. SNF1-RELATED KINASE 1 and TARGET OF RAPAMYCIN control light-responsive splicing events and developmental characteristics in etiolated Arabidopsis seedlings. Plant Cell 35, 3413–3428; 10.1093/plcell/koad168 (2023).

69. Stuttmann, J. et al. Highly efficient multiplex editing: one-shot generation of 8× Nicotiana benthamiana and 12× Arabidopsis mutants. Plant J 106, 8–22; 10.1111/tpj.15197 (2021).

70. Liu, H. et al. CRISPR-P 2.0: An Improved CRISPR-Cas9 Tool for Genome Editing in Plants. Mol Plant 10, 530–532; 10.1016/j.molp.2017.01.003 (2017).

71. Patro, R., Duggal, G., Love, M. I., Irizarry, R. A. & Kingsford, C. Salmon provides fast and bias-aware quantification of transcript expression. Nat Methods 14, 417–419; 10.1038/nmeth.4197 (2017).

72. Guo, W. et al. 3D RNA-seq: a powerful and flexible tool for rapid and accurate differential expression and alternative splicing analysis of RNA-seq data for biologists. RNA Biol 18, 1574–1587; 10.1080/15476286.2020.1858253 (2021).

73. Sherman, B. T. et al. DAVID: a web server for functional enrichment analysis and functional annotation of gene lists (2021 update). Nucleic acids research 50, W216–W221; 10.1093/nar/gkac194 (2022).

