## Supplemental Figures for "A structured RNA balances DEAD-box RNA helicase function in plant alternative splicing control"

#### Supplemental material

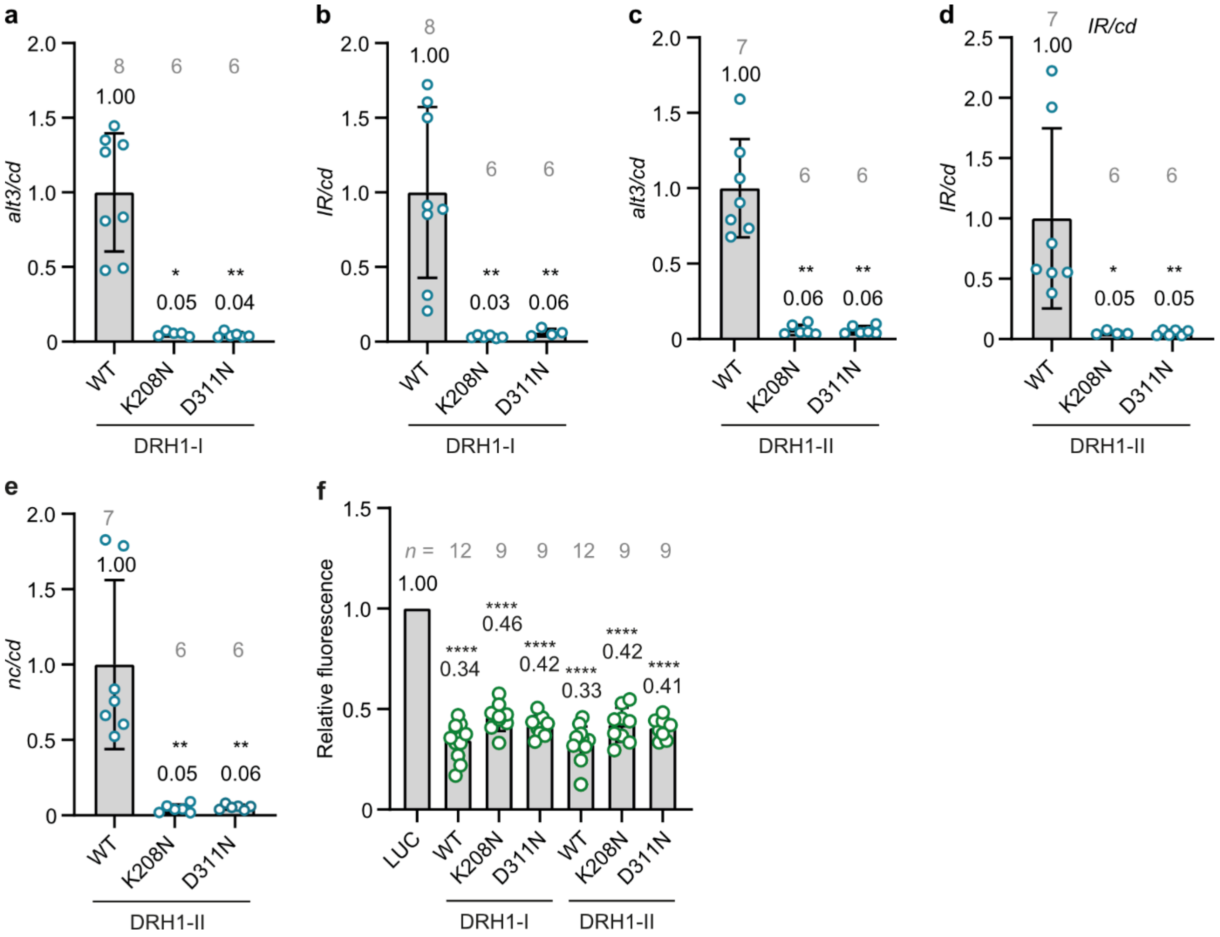


**Supplemental Figure 1: Helicase activity of both DRH1 isoforms is required for autoregulation via AS. a-e:** AS of *DRH1* reporter upon transient expression in *N. benthamiana* when co-transformed with WT DRH1-I, WT DRH1-II or their respective mutants impaired in helicase activity. AS ratio of reporter with WT DRH1-I (**a-b**) or WT DRH1-II (**c-e**) was set to 1. *n* is provided in grey letters above each bar, missing data points are due to undetectable levels of the respective AS variant in individual samples. **f:** DRH1 reporter fluorescence upon transient expression in *N. benthamiana* when co-transformed with WT DRH1-I, WT DRH1-II or their respective mutants impaired in helicase activity. *n* is provided in grey letters above each bar. Mean values, standard deviations; individual data points represent biological replicates and are depicted as colored dots. Asterisks indicate significant change compared to co-transformation of WT DRH1 (**a, c-e**: Kruskal-Wallis test followed by Dunn's multiple comparisons test, **b:** Brown-Forsythe and Welch ANOVA followed by Dunnett's T3 multiple comparisons test) or LUC (**f**, one-sample t-test). *p < 0.05, **p < 0.01, ****p < 0.0001.

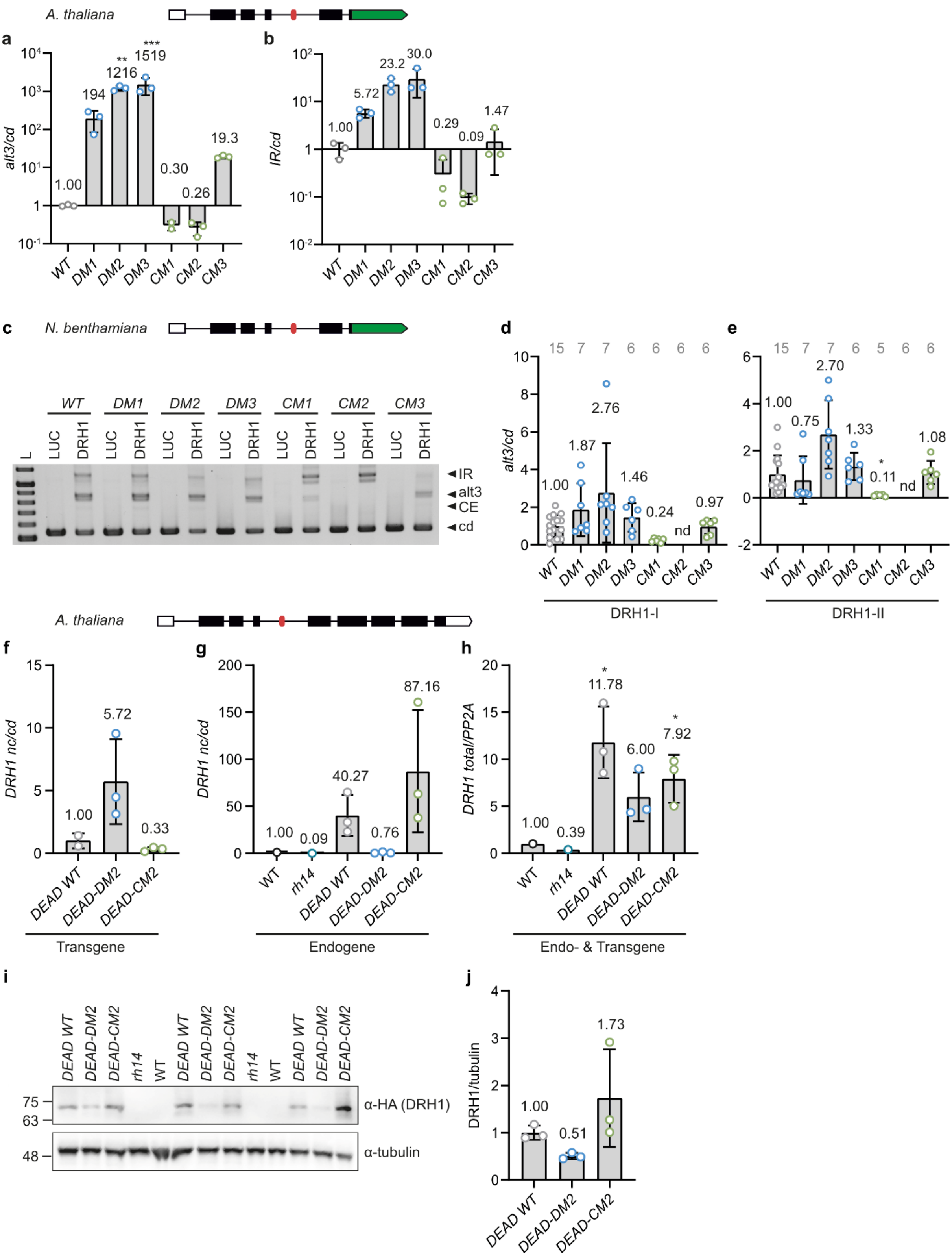
A. thaliana
a
alt3/cd
1.00
194
**
1216
***
1519
0.30
0.26
19.3
WT DM1 DM2 DM3 CM1 CM2 CM3
b
IR/cd
1.00
5.72
23.2
30.0
0.29
0.09
1.47
WT DM1 DM2 DM3 CM1 CM2 CM3
c
N. benthamiana
WT DM1 DM2 DM3 CM1 CM2 CM3
L
LUC DRH1
◄IR
◄alt3
◄CE
◄cd
d
15 7 7 6 6 6 6
alt3/cd
1.00
1.87
2.76
1.46
0.24
nd
0.97
WT DM1 DM2 DM3 CM1 CM2 CM3
DRH1-I
e
15 7 7 6 5 6 6
1.00
0.75
2.70
1.33
*
0.11
nd
1.08
WT DM1 DM2 DM3 CM1 CM2 CM3
DRH1-II
A. thaliana
f
DRH1 nc/cd
1.00
5.72
0.33
DEAD WT DEAD-DM2 DEAD-CM2
Transgene
g
DRH1 nc/cd
1.00
0.09
40.27
0.76
87.16
WT rh14 DEAD WT DEAD-DM2 DEAD-CM2
Endogene
h
DRH1 total/PP2A
1.00
0.39
*
11.78
6.00
*
7.92
WT rh14 DEAD WT DEAD-DM2 DEAD-CM2
Endo- & Transgene
i
DEAD WT DEAD-DM2 DEAD-CM2 rh14 WT DEAD WT DEAD-DM2 DEAD-CM2 rh14 WT DEAD WT DEAD-DM2 DEAD-CM2
75
63
48
α-HA (DRH1)
α-tubulin
j
DRH1/tubulin
1.00
0.51
1.73
DEAD WT DEAD-DM2 DEAD-CM2

**Supplemental Figure 2: *DEAD*-mediated autoregulation of *DRH1*. a-b:** Bioanalyzer quantification of WT and *DEAD* mutant reporter AS in stably transformed *A. thaliana* lines, AS ratio of WT reporter was set to 1. $n = 3$, missing data points are due to undetectable levels of the respective AS variant in individual samples. Asterisks indicate significant change compared to the WT reporter (**a:** one-way ANOVA followed by Dunnett's multiple comparisons test, **b:** Kruskal-Wallis test followed by Dunn's multiple comparisons test. **$p < 0.01$, ***$p < 0.001$). **c:** Representative gel image of *DRH1* WT and *DEAD* mutant reporter AS in *N. benthamiana* co-transformed with DRH1-**I**. L: ladder, from 0.5-1 kb in 0.1 kb increments, 1.2 kb. **d-e:** Bioanalyzer quantification of *DRH1* WT and *DEAD* mutant reporter AS upon transient expression in *N. benthamiana* when DRH1-**I** (**d**) or DRH1-**II** (**e**) is co-transformed. AS ratio of the WT reporter with the respective isoform is set to 1, nd: not detectable (complete absence of *alt3* variant). *n* is given in grey letters above each bar. Asterisk indicates significant change compared to the WT reporter (Kruskal-Wallis test followed by Dunn's multiple comparisons test, *$p < 0.05$). **f-g:** Bioanalyzer quantification of the AS pattern of *pUBQ:DRH1* genomic constructs carrying different mutations of *DEAD* in *rh14* background (**f**) or endogenous *DRH1* in the same plants (**g**). Splicing ratio of the *pUBQ:DRH1* construct with WT *DEAD* motif (**f**) or of WT plants (**g**) was set to 1. $n = 1$ for WT and *rh14* controls, $n = 3$ for *pUBQ:DRH1* lines. For *DEAD WT* construct in **f**, one sample was excluded due to inconsistent marker peaks in Bioanalyzer run. No significant change was detected compared to WT *DEAD* motif according to one-way ANOVA followed by Dunnett's multiple comparisons test. **h:** qPCR quantification of total *DRH1* transcript levels in the same mutants as in **f** and **g**. Transcript levels in WT plants were set to 1. $n = 1$ for WT and *rh14* controls, $n = 3$ for *pUBQ:DRH1* lines. Asterisks indicate significant change compared to WT background (one-sample t-test, *$p < 0.05$). **i-j:** Western blot (**i**) and quantification of grey values (**j**) for the same mutants as in **f-h**. 3 independent lines per genotype were used, mean grey values for DRH1 and tubulin signals, respectively, were determined using ImageJ. Mean values, standard deviations; individual data points represent biological replicates (**d-e**) or independent lines (**a-b, f-h, j**) and are depicted as colored dots.

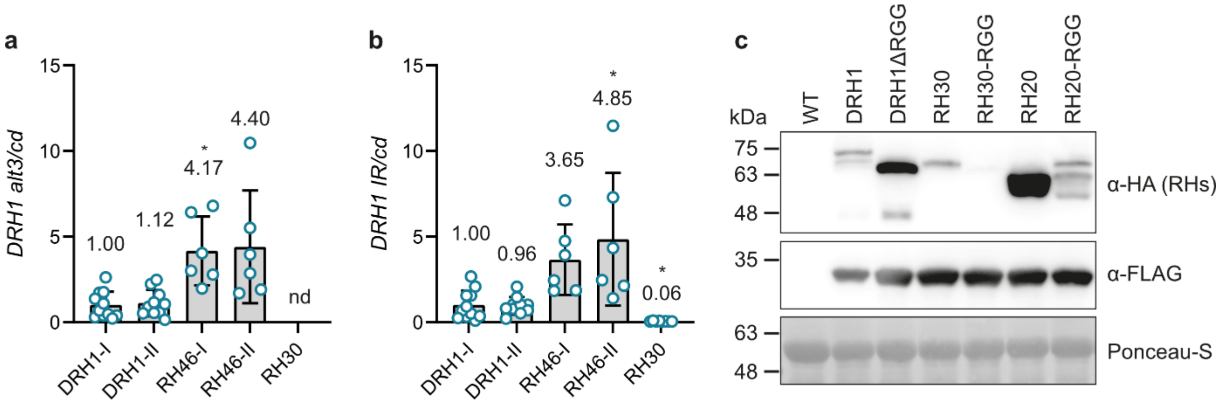


**Supplemental Figure 3: RH46 and RH30 can crossregulate *DRH1* via AS in *N. benthamiana*. a-b:** Bioanalyzer quantification of *DRH1* reporter AS on transcript level upon transient co-expression of DRH1, RH46, and RH30. AS ratio in presence of DRH1-**I** was set to 1, nd: not detectable. $n$ = 11 for DRH1-**I**, 10 for DRH1-**II**, 6 for both RH46 isoforms, and 8 for RH30. Mean values, standard deviations; individual data points represent biological replicates and are depicted as colored dots. Asterisks indicate significant change compared to DRH1-**I** (**a:** Brown-Forsythe and Welch ANOVA followed by Dunnett's T3 multiple comparisons test, **b:** Kruskal-Wallis test followed by Dunn's multiple comparisons test, $^*p < 0.05$). **c:** Western Blot showing transient expression of RGG/RG motif mutant helicases in *N. benthamiana*. Detection of helicases (α-HA), co-expressed mOrange2 (α-FLAG) or total protein (Ponceau-S). Expected sizes: DRH1, 69 kDa; DRH1ΔRGG, 60 kDa; RH30, 66 kDa; RH30-RGG, 70 kDa; RH20, 57 kDa; RH20-RGG, 63 kDa.

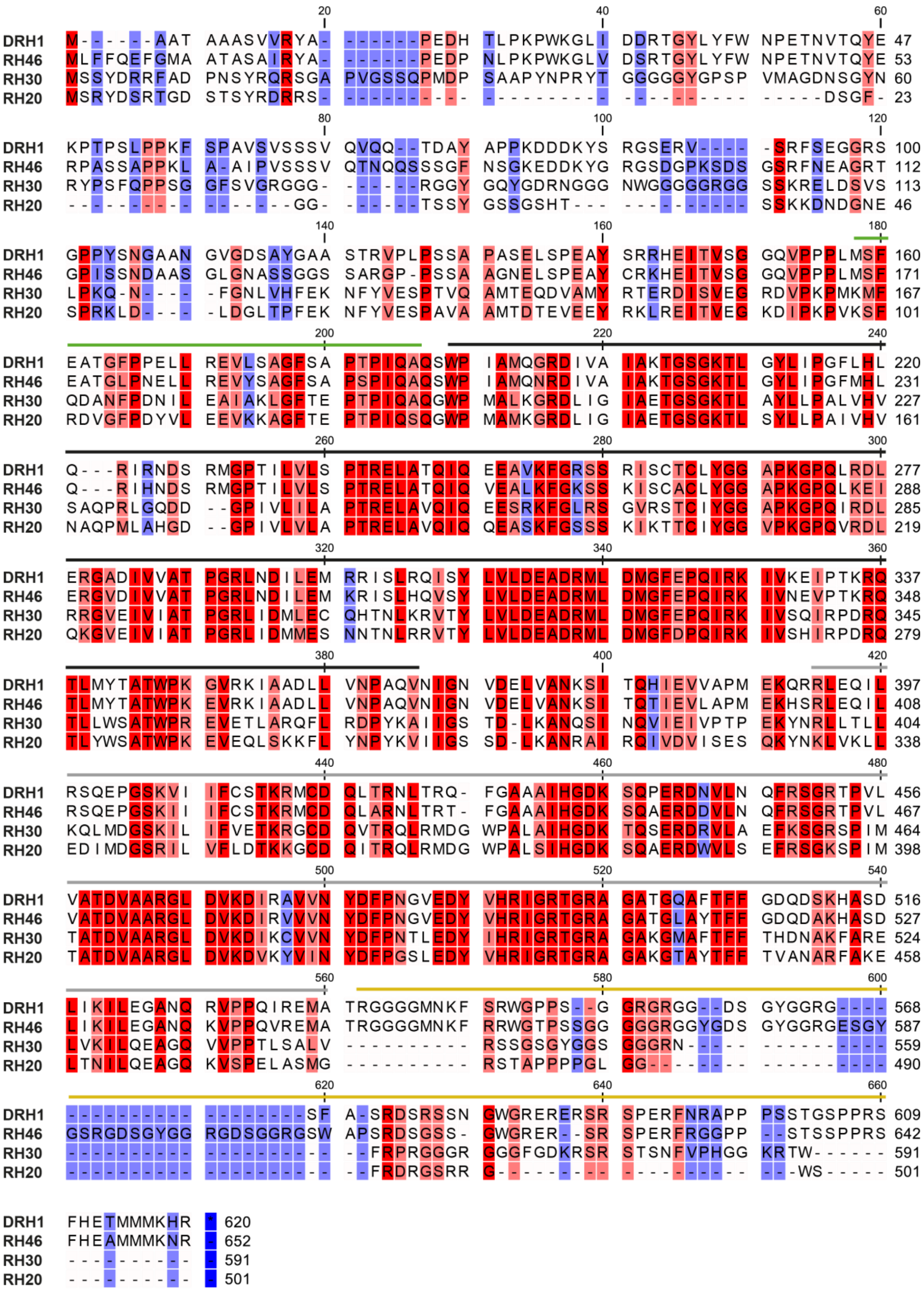
DRH1 M------AAT AAASVVRYA- ------PEDH TLPKPWKGLI DDRTGYLYFW NPETNVTQYE 47
RH46 MLFFQEFGMA ATASAIRYA- ------PEDP NLPKPWKGLV DSRTGYLYFW NPETNVTQYE 53
RH30 MSSYDRRFAD PNSYRQRSGA PVGSSQPMDP SAAPYNPRYT GGGGGYGPSP VMAGDNSGYN 60
RH20 MSRYDSRTGD STSYRDRRS- ---------- ---------- ---------- -----DSGF- 23
DRH1 KPTPSLPPKF SPAVSVSSSV QVQQ--TDAY APPKDDDKYS RGSERV---- -SRFSEGGRS 100
RH46 RPASSAPPKL A-AIPVSSSV QTNQQSSSGF NSGKEDDKYG RGSDGPKSDS GSRFNEAGRT 112
RH30 RYPSFQPPSG GFSVGRGGG- ------RGGY GQYGDRNGGG NWGGGGGRGG SSKRELDSVS 113
RH20 ---------- -------GG- ------TSSY GSSGSHT--- ---------- SSKKDNDGNE 46
DRH1 GPPYSNGAAN GVGDSAYGAA STRVPLPSSA PASELSPEAY SRRHEITVSG GQVPPPLMSF 160
RH46 GPISSNDAAS GLGNASSGGS SARGP-PSSA AGNELSPEAY CRKHEITVSG GQVPPPLMSF 171
RH30 LPKQ-N---- -FGNLVHFEK NFYVESPTVQ AMTEQDVAMY RTERDISVEG RDVPKPMKMF 167
RH20 SPRKLD---- -LDGLTPFEK NFYVESPAVA AMTDTEVEEY RKLREITVEG KDIPKPVKSF 101
DRH1 EATGFPPELL REVLSAGFSA PTPIQAQSWP IAMQGRDIVA IAKTGSGKTL GYLIPGFLHL 220
RH46 EATGLPNELL REVYSAGFSA PSPIQAQSWP IAMQNRDIVA IAKTGSGKTL GYLIPGFMHL 231
RH30 QDANFPDNIL EAIAKLGFTE PTPIQAQGWP MALKGRDLIG IAETGSGKTL AYLLPALVHV 227
RH20 RDVGFPDYVL EEVKKAGFTE PTPIQSQGWP MAMKGRDLIG IAETGSGKTL SYLLPAIVHV 161
DRH1 Q---RIRNDS RMGPTILVLS PTRELATQIQ EEAVKFGRSS RISCTCLYGG APKGPQLRDL 277
RH46 Q---RIHNDS RMGPTILVLS PTRELATQIQ VEALKFGKSS KISCACLYGG APKGPQLKEI 288
RH30 SAQPRLGQDD --GPIVLILA PTRELAVQIQ EESRKFGLRS GVRSTCIYGG APKGPQIRDL 285
RH20 NAQPMLAHGD --GPIVLVLA PTRELAVQIQ QEASKFGSSS KIKTTCIYGG VPKGPQVRDL 219
DRH1 ERGADIVVAT PGRLNDILEM RRISLRQISY LVLDEADRML DMGFEPQIRK IVKEIPTKRQ 337
RH46 ERGVDIVVAT PGRLNDILEM KRISLHQVSY LVLDEADRML DMGFEPQIRK IVNEVPTKRQ 348
RH30 RRGVEIVIAT PGRLIDMLEC QHTNLKRVTY LVLDEADRML DMGFEPQIRK IVSQIRPDRQ 345
RH20 QKGVEIVIAT PGRLIDMMES NNTNLRRVTY LVLDEADRML DMGFDPQIRK IVSHIRPDRQ 279
DRH1 TLMYTATWPK GVRKIAADLL VNPAQVNIGN VDELVANKSI TQHIEVVAPM EKQRRLEQIL 397
RH46 TLMYTATWPK EVRKIAADLL VNPAQVNIGN VDELVANKSI TQTIEVLAPM EKHSRLEQIL 408
RH30 TLLWSATWPR EVETLARQFL RDPYKAIIGS TD-LKANQSI NQVIEIVPTP EKYNRLTLLL 404
RH20 TLYWSATWPK EVEQLSKKFL YNPYKVIIGS SD-LKANRAI RQIVDVISES QKYNKLVKLL 338
DRH1 RSQEPGSKVI IFCSTKRMCD QLTRNLTRQ- FGAAAIHGDK SQPERDNVLN QFRSGRTPVL 456
RH46 RSQEPGSKII IFCSTKRMCD QLARNLTRT- FGAAAIHGDK SQAERDDVLN QFRSGRTPVL 467
RH30 KQLMDGSKIL IFVETKRGCD QVTRQLRMDG WPALAIHGDK TQSERDRVLA EFKSGRSPIM 464
RH20 EDIMDGSRIL VFLDTKKGCD QITRQLRMDG WPALSIHGDK SQAERDWVLS EFRSGKSPIM 398
DRH1 VATDVAARGL DVKDIRAVVN YDFPNGVEDY VHRIGRTGRA GATGQAFTFF GDQDSKHASD 516
RH46 VATDVAARGL DVKDIRVVVN YDFPNGVEDY VHRIGRTGRA GATGLAYTFF GDQDAKHASD 527
RH30 TATDVAARGL DVKDIKCVVN YDFPNTLEDY IHRIGRTGRA GAKGMAFTFF THDNAKFARE 524
RH20 TATDVAARGL DVKDVKYVIN YDFPGSLEDY VHRIGRTGRA GAKGTAYTFF TVANARFAKE 458
DRH1 LIKILEGANQ RVPPQIREMA TRGGGGMNKF SRWGPPS--G GRGRGG--DS GYGGRG---- 568
RH46 LIKILEGANQ KVPPQVREMA TRGGGGMNKF RRWGTPSSGG GGGRGGYGDS GYGGRGESGY 587
RH30 LVKILQEAGQ VVPPTLSALV ---------- RSSGSGYGGS GGGRN----- ---------- 559
RH20 LTNILQEAGQ KVSPELASMG ---------- RSTAPPPPGL GG-------- ---------- 490
DRH1 ---------- --------SF A-SRDSRSSN GWGRERERSR SPERFNRAPP PSSTGSPPRS 609
RH46 GSRGDSGYGG RGDSGGRGSW APSRDSGSS- GWGRER--SR SPERFRGGPP --STSSPPRS 642
RH30 ---------- ---------- --FRPRGGGR GGGFGDKRSR STSNFVPHGG KRTW------ 591
RH20 ---------- ---------- --FRDRGSRR G--------- ---------- ---WS----- 501
DRH1 FHETMMMKHR * 620
RH46 FHEAMMMKNR - 652
RH30 ---------- - 591
RH20 ---------- - 501

**Supplemental Figure 4: Alignment of the amino acid sequences of DRH1, RH46, RH30 and RH20.** Blue and red shading indicates unconserved and conserved residues, respectively; with darker shades of red pointing out higher degrees of conservation. Location of conserved domains was inferred from www.uniprot.org and is shown by coloured lines above the alignment (from N- to C-terminus: Q-motif (green), N-terminal helicase domain (black), C-terminal helicase domain (grey), RGG box (yellow)). Note that the RGG box is only present in DRH1 and RH46. Numbers at the right indicate the amino acid position in the respective protein. Alignment was created with CLC Main Workbench 20.0.4 (Qiagen).

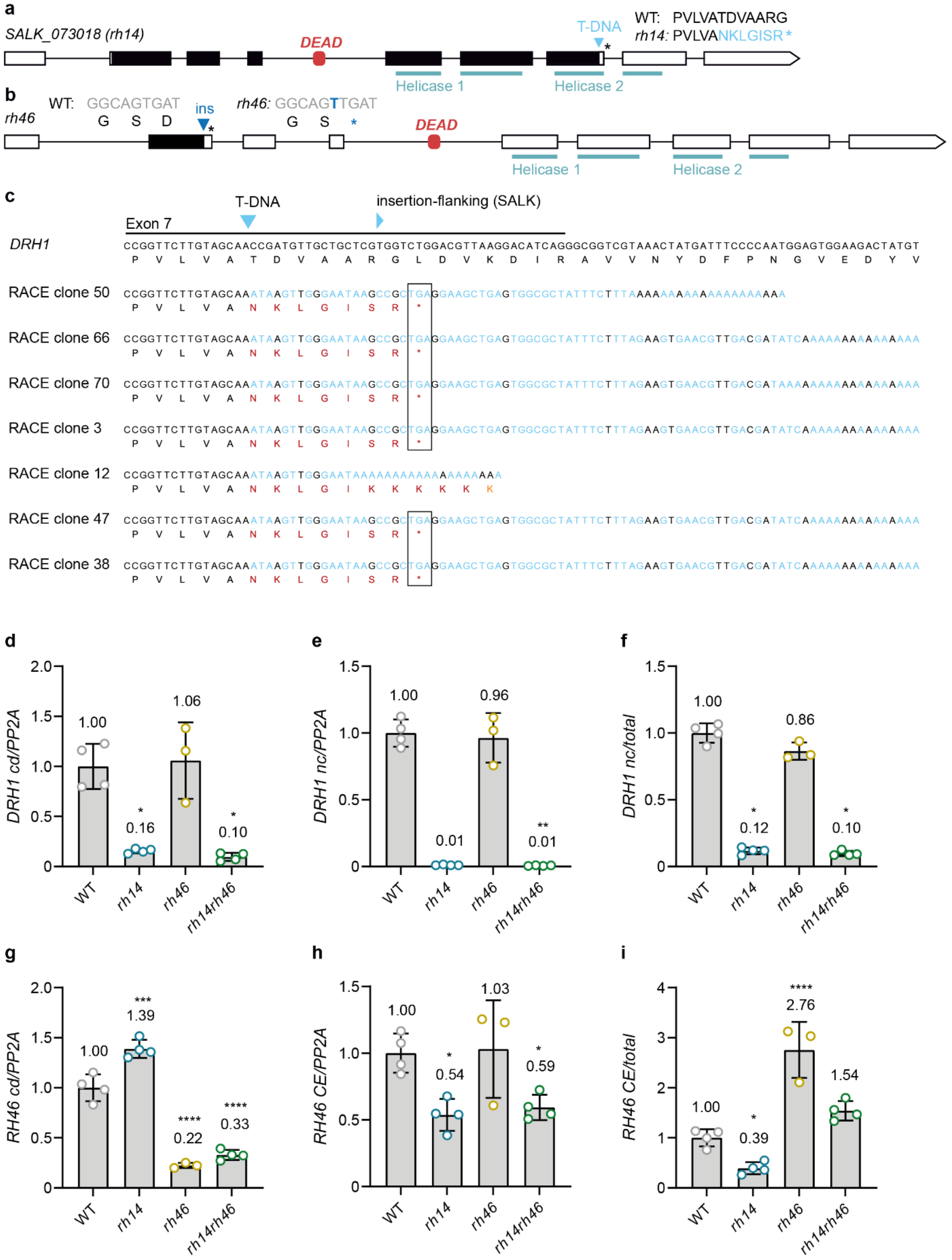

a
SALK_073018 (rh14)
DEAD
T-DNA
WT: PVLVATDVAARG
rh14: PVLVANKLGISR*
Helicase 1
Helicase 2
b
rh46
WT: GGCAGTGAT
G S D
ins
rh46: GGCAGTTGAT
G S *
DEAD
Helicase 1
Helicase 2
c
T-DNA
insertion-flanking (SALK)
Exon 7
DRH1
CCGGTTCTTGTAGCAACCGATGTTGCTGCTCGTGGTCTGGACGTTAAGGACATCAGGGCGGTCGTAAACTATGATTTCCCCAATGGAGTGGAAGACTATGT
P V L V A T D V A A R G L D V K D I R A V V N Y D F P N G V E D Y V
RACE clone 50
CCGGTTCTTGTAGCAAATAAGTTGGGAATAAGCCGCTGAGGAAGCTGAGTGGCGCTATTTCTTTAAAAAAAAAAAAAAAAAAAAA
P V L V A N K L G I S R *
RACE clone 66
CCGGTTCTTGTAGCAAATAAGTTGGGAATAAGCCGCTGAGGAAGCTGAGTGGCGCTATTTCTTTAGAAGTGAACGTTGACGATATCAAAAAAAAAAAAAAAA
P V L V A N K L G I S R *
RACE clone 70
CCGGTTCTTGTAGCAAATAAGTTGGGAATAAGCCGCTGAGGAAGCTGAGTGGCGCTATTTCTTTAGAAGTGAACGTTGACGATAAAAAAAAAAAAAAAAAAA
P V L V A N K L G I S R *
RACE clone 3
CCGGTTCTTGTAGCAAATAAGTTGGGAATAAGCCGCTGAGGAAGCTGAGTGGCGCTATTTCTTTAGAAGTGAACGTTGACGATATCAAAAAAAAAAAAAAAA
P V L V A N K L G I S R *
RACE clone 12
CCGGTTCTTGTAGCAAATAAGTTGGGAATAAAAAAAAAAAAAAAAAAAA
P V L V A N K L G I K K K K K K
RACE clone 47
CCGGTTCTTGTAGCAAATAAGTTGGGAATAAGCCGCTGAGGAAGCTGAGTGGCGCTATTTCTTTAGAAGTGAACGTTGACGATATCAAAAAAAAAAAAAAAA
P V L V A N K L G I S R *
RACE clone 38
CCGGTTCTTGTAGCAAATAAGTTGGGAATAAGCCGCTGAGGAAGCTGAGTGGCGCTATTTCTTTAGAAGTGAACGTTGACGATATCAAAAAAAAAAAAAAAA
P V L V A N K L G I S R *
d
DRH1 cd/PP2A
2.0
1.5
1.0
0.5
0
1.00
*
0.16
1.06
*
0.10
WT
rh14
rh46
rh14rh46
e
DRH1 nc/PP2A
1.5
1.0
0.5
0
1.00
0.01
0.96
**
0.01
WT
rh14
rh46
rh14rh46
f
DRH1 nc/total
1.5
1.0
0.5
0
1.00
*
0.12
0.86
*
0.10
WT
rh14
rh46
rh14rh46
g
RH46 cd/PP2A
2.0
1.5
1.0
0.5
0
1.00
***
1.39
****
0.22
****
0.33
WT
rh14
rh46
rh14rh46
h
RH46 CE/PP2A
1.5
1.0
0.5
0
1.00
*
0.54
1.03
*
0.59
WT
rh14
rh46
rh14rh46
i
RH46 CE/total
4
3
2
1
0
1.00
*
0.39
****
2.76
1.54
WT
rh14
rh46
rh14rh46

**Supplemental Figure 5: Auto- and crossregulation of *DRH1* and *RH46*. a-b:** Gene models depicting the mutations in *rh14* (**a**) and *rh46* (**b**) plants. Exons, introns, CDS and UTRs are depicted as boxes, lines, black and white shading, respectively; *DEAD* motif is shown as red rounded rectangle. Asterisks indicate stop codons, mutations (*rh14*: T-DNA insertion, *rh46*: one-nucleotide insertion due to CRISPR-Cas9 mutagenesis) are illustrated as blue triangles. Location of the helicase domains was inferred from www.uniprot.org and is shown with turquoise lines. The amino acid (**a**) or nucleotide and amino acid sequences (**b**) in WT and mutant background are given above the respective gene model in grey and black letters, respectively; mutations are depicted in blue. **c:** Alignment of *DRH1* WT sequence and different clones obtained by 3' RACE of *rh14* plants. Blue triangles indicate start of insertion-flanking sequence according to SALK and actual position of the T-DNA. Nucleotides and amino acids not corresponding to the WT sequence are shown in blue and red, respectively. Stop codon used for the constructs in **Fig. 5a-c** is indicated by a black box. **d-i:** Quantitative PCR of *DRH1* (**d-f**) and *RH46* (**g-i**) transcripts in *A. thaliana* seedlings of different backgrounds. Data points represent biological replicates, transcript ratio of WT seedlings is set to 1. $n$ = 3 for *rh46*, $n$ = 4 for all others. Mean values, standard deviations; individual data points are depicted as colored dots. Asterisks indicate significant change compared to WT background (**d:** Brown-Forsythe and Welch ANOVA followed by Dunnett's T3 multiple comparisons test, **e-f:** Kruskal-Wallis test followed by Dunn's multiple comparisons test, **g-i:** one-way ANOVA followed by Dunnett's multiple comparisons test, *$p < 0.05$, **$p < 0.01$, ****$p < 0.0001$).

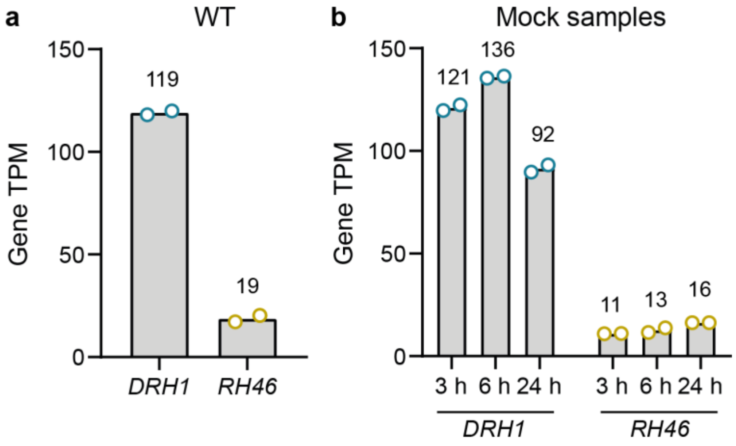


**Supplemental Figure 6: *DRH1* is more highly expressed than *RH46* in seedlings.** Transcripts per kilobase million (TPM) values for *DRH1* and *RH46* in WT seedlings (**a**) and seedlings of inducible DRH1 overexpression lines treated with mock solution for different time intervals (**b**). Bars show mean values ($n$ = 2), individual data points represent biological replicates and are depicted as colored dots.

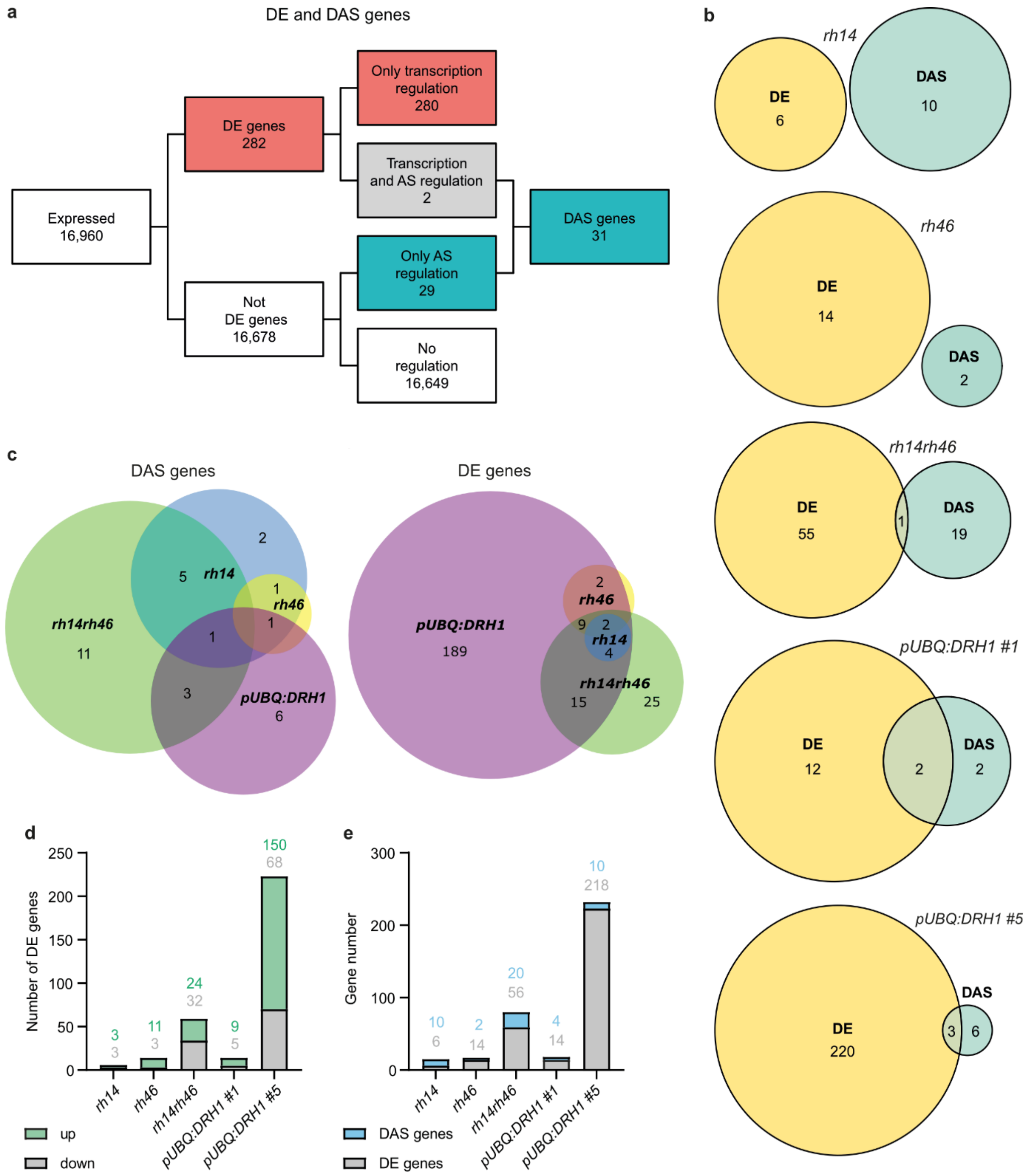


**Supplemental Figure 7: Knockout of *DRH1* and/or *RH46* has little effect on the transcriptome. a:** Overview of the numbers of expressed, differentially expressed (DE), and differentially alternatively spliced (DAS) genes in at least one mutant compared to the WT, using the following cut-offs: P-value < 0.01; log2 fold change (DE) > 1; deltaPS (DAS) > 0.1. **b:** Venn diagrams showing the numbers and overlaps between significant (p < 0.01) DE and DAS targets in the individual mutants compared to the WT. **c:** Overlap between the targets of different mutant lines. Targets of the two *pUBQ:DRH1* lines were combined. Diagrams were created with DeepVenn 1. **d:** Numbers of DE genes that are up- or downregulated in the individual mutants compared to the WT. **e:** Total numbers of DE and DAS genes in the individual mutants compared to the WT.

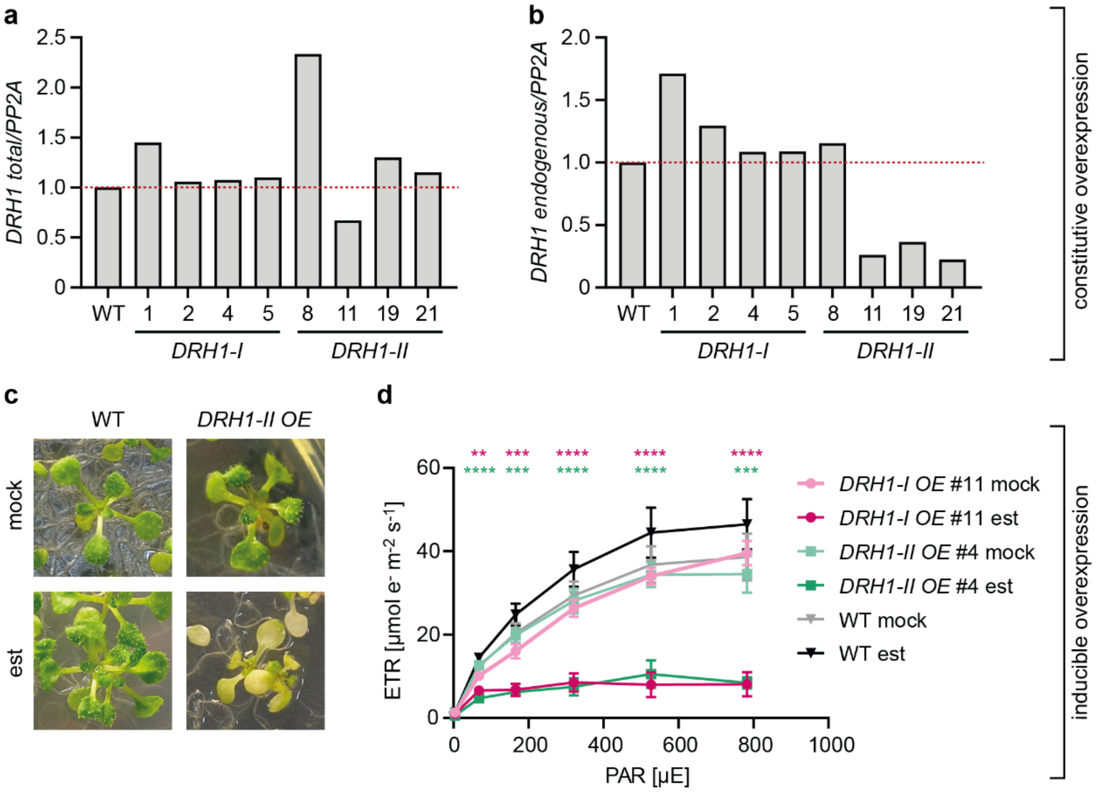


**Supplemental Figure 8: Both isoforms of DRH1 cause a stress phenotype when overexpressed. a-b:** Quantitative PCR of total *DRH1* (**a**) and total endogenous *DRH1* transcript levels (**b**) in seedlings constitutively expressing *DRH1-I* or *DRH1-II* CDS constructs. Each bar corresponds to an independent line that was measured in a single replicate. Transcript levels in WT seedlings were set to 1 and are indicated by the red dotted line. **c:** 14-day-old seedlings (WT or the same *DRH1-II* overexpression line as in **Fig. 6c**) 4 days after mock or estradiol treatment. **d:** Electron transport rate (ETR) calculated as ETR = Y(II)*PAR*0.5*0.84 in WT seedlings or the same two *DRH1* overexpression lines as in **c** and **Fig. 6b-e** 4 days after induction. The same seedlings were measured at multiple light intensities. The data for WT and *DRH1-I OE* #11 is the same as in **Fig. 6e**. PAR: photosynthetically active radiation, mean values ($n$ = 5), standard deviations. Asterisks indicate significant change between estradiol and mock treatment (two-way repeated measures ANOVA followed by Šídák's multiple comparisons test; **$p < 0.01$, ***$p < 0.001$, ****$p < 0.0001$).

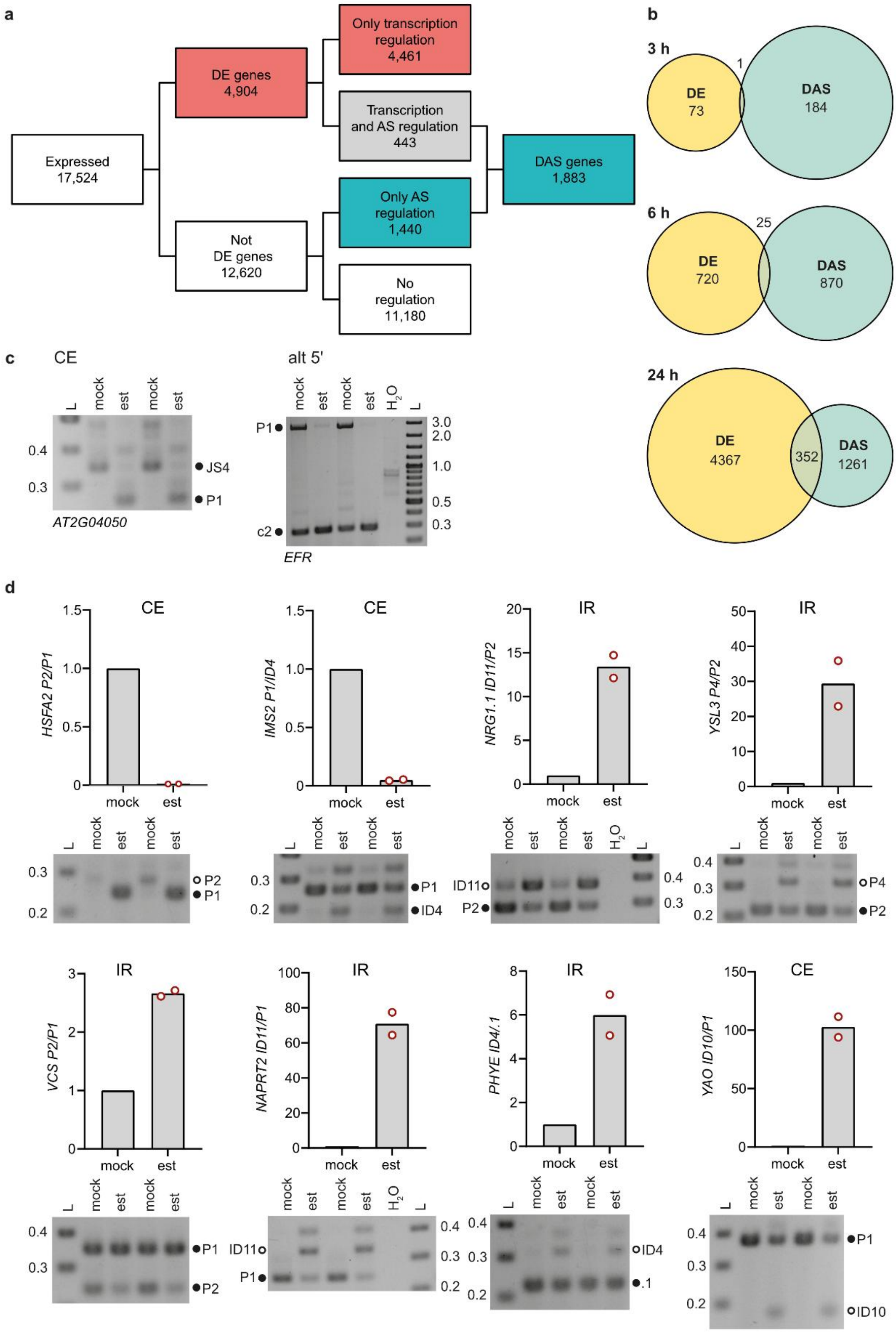
a
Expressed
17,524
DE genes
4,904
Not
DE genes
12,620
Only transcription
regulation
4,461
Transcription
and AS regulation
443
Only AS
regulation
1,440
No
regulation
11,180
DAS genes
1,883
b
3 h
DE
73
1
DAS
184
6 h
DE
720
25
DAS
870
24 h
DE
4367
352
DAS
1261
c
CE
L
mock
est
mock
est
0.4
0.3
JS4
P1
AT2G04050
alt 5'
mock
est
mock
est
H2O
L
P1
c2
3.0
2.0
1.0
0.5
0.3
EFR
d
CE
HSFA2 P2/P1
mock
est
P2
P1
CE
IMS2 P1/ID4
mock
est
P1
ID4
IR
NRG1.1 ID11/P2
mock
est
ID11
P2
IR
YSL3 P4/P2
mock
est
P4
P2
IR
VCS P2/P1
mock
est
P1
P2
IR
NAPRT2 ID11/P1
mock
est
ID11
P1
IR
PHYE ID4/.1
mock
est
ID4
.1
CE
YAO ID10/P1
mock
est
P1
ID10

**Supplemental Figure 9: Validation of RNA-seq results from inducible overexpression lines. a:** Overview of the numbers of expressed, differentially expressed (DE), and differentially alternatively spliced (DAS) genes for at least one time point after induction of *DRH1* overexpression. **b:** Venn diagrams showing the numbers and overlaps between significant ($p < 0.01$) DE and DAS targets 3 h, 6 h, and 24 h after induction of *DRH1* overexpression. **c-d:** Co-amplification PCRs of significant DAS genes for two replicates each 6 h after estradiol or mock treatment. Bioanalyzer quantification is given at the top, representative gel image at the bottom. Diagram titles indicate the type of AS event, splicing ratios are normalized to respective mock samples. For the two events in **c**, no quantification was possible either due to complete splicing to one variant under mock conditions (*AT2G04050*) or due to fragment size exceeding Bioanalyzer specifications (*EFR).* L: ladder, band sizes are given in kb. Relevant transcript variants are marked with dots, naming corresponds to AtRTD2-QUASI 2. Variants introducing a PTC are represented by white dots, potentially coding ones are marked in black. Bars show mean values ($n$ = 2), individual data points represent biological replicates and are shown as red dots.

### References


1. Hulsen, T. DeepVenn -- a web application for the creation of area-proportional Venn diagrams using the deep learning framework Tensorflow.js, 9/27/2022.

2. Zhang, R. *et al.* A high quality Arabidopsis transcriptome for accurate transcript-level analysis of alternative splicing. *Nucleic acids research* **45,** 5061–5073; 10.1093/nar/gkx267 (2017).
